# Evolution of Antibody Immunity to SARS-CoV-2

**DOI:** 10.1101/2020.11.03.367391

**Authors:** Christian Gaebler, Zijun Wang, Julio C. C. Lorenzi, Frauke Muecksch, Shlomo Finkin, Minami Tokuyama, Alice Cho, Mila Jankovic, Dennis Schaefer-Babajew, Thiago Y. Oliveira, Melissa Cipolla, Charlotte Viant, Christopher O. Barnes, Yaron Bram, Gaëlle Breton, Thomas Hägglöf, Pilar Mendoza, Arlene Hurley, Martina Turroja, Kristie Gordon, Katrina G. Millard, Victor Ramos, Fabian Schmidt, Yiska Weisblum, Divya Jha, Michael Tankelevich, Gustavo Martinez-Delgado, Jim Yee, Roshni Patel, Juan Dizon, Cecille Unson-O’Brien, Irina Shimeliovich, Davide F. Robbiani, Zhen Zhao, Anna Gazumyan, Robert E. Schwartz, Theodora Hatziioannou, Pamela J. Bjorkman, Saurabh Mehandru, Paul D. Bieniasz, Marina Caskey, Michel C. Nussenzweig

**Affiliations:** Laboratory of Molecular Immunology, The Rockefeller University, New York, NY 10065, USA; Laboratory of Retrovirology, The Rockefeller University, New York, NY 10065, USA; Division of Gastroenterology, Department of Medicine, Icahn School of Medicine at Mount Sinai, New York, NY 10029, USA; Division of Biology and Biological Engineering, California Institute of Technology, Pasadena, CA, USA; Division of Gastroenterology and Hepatology, Department of Medicine, Weill Cornell Medicine, New York, NY 10065, USA; Hospital Program Direction, The Rockefeller University, New York, NY 10065, USA; Department of Pathology and Laboratory Medicine, Weill Cornell Medicine, New York, NY 10065, USA; Institute for Research in Biomedicine, Università della Svizzera italiana, Bellinzona, Switzerland; Department of Physiology, Biophysics and Systems Biology, Weill Cornell Medicine, New York, NY 10065, USA; Howard Hughes Medical Institute

## Abstract

Severe acute respiratory syndrome coronavirus-2 (SARS-CoV-2) has infected 78 million individuals and is responsible for over 1.7 million deaths to date. Infection is associated with development of variable levels of antibodies with neutralizing activity that can protect against infection in animal models. Antibody levels decrease with time, but the nature and quality of the memory B cells that would be called upon to produce antibodies upon re-infection has not been examined. Here we report on the humoral memory response in a cohort of 87 individuals assessed at 1.3 and 6.2 months after infection. We find that IgM, and IgG anti-SARS-CoV-2 spike protein receptor binding domain (RBD) antibody titers decrease significantly with IgA being less affected. Concurrently, neutralizing activity in plasma decreases by five-fold in pseudotype virus assays. In contrast, the number of RBD-specific memory B cells is unchanged. Memory B cells display clonal turnover after 6.2 months, and the antibodies they express have greater somatic hypermutation, increased potency and resistance to RBD mutations, indicative of continued evolution of the humoral response. Analysis of intestinal biopsies obtained from asymptomatic individuals 4 months after coronavirus disease-2019 (COVID-19) onset, using immunofluorescence, or polymerase chain reaction, revealed persistence of SARS-CoV-2 nucleic acids and immunoreactivity in the small bowel of 7 out of 14 volunteers. We conclude that the memory B cell response to SARS-CoV-2 evolves between 1.3 and 6.2 months after infection in a manner that is consistent with antigen persistence.

---

Antibody responses to SARS-CoV-2 were initially characterized in a cohort of COVID-19-convalescent individuals approximately 40 days (1.3 months) after infection ^1^. Between 31 August and 16 October 2020, 100 participants returned for a 6-month follow-up study visit. Although initial criteria allowed enrollment of close contacts of individuals diagnosed with reverse transcription (RT–PCR) confirmed SARS-CoV-2 infection ^1^, 13 of the contacts did not seroconvert and were excluded from further analyses. The remaining 87 participants with RT-PCR–confirmed COVID-19 diagnosis and/or seroconversion returned for analysis approximately 191 days (6.2 months, range: 165-223 days) after the onset of symptoms. In this cohort, symptoms lasted for a median of 12 days (0–44 days) during the acute phase, and 10 (11%) of the participants were hospitalized. Consistent with other reports ^2,3^, 38 (44%) of the participants reported persistent long-term symptoms attributable to COVID-19 (Methods and Supplementary Tables 1 and 2). The duration and severity of symptoms during acute disease was significantly greater among participants with persistent post-acute symptoms at the second study visit (Extended Data Fig. 1m-q). Importantly, all 87 participants tested negative for SARS-CoV-2 at the 6-month follow-up study visit using an approved saliva-based PCR assay (Methods). Participant demographics and clinical characteristics are shown in Supplementary Tables 1,2.

Antibody reactivity in plasma to RBD and nucleoprotein (N) was measured by enzyme-linked immunosorbent assay (ELISA) and automated serological assays ^1,4,5^. Anti-RBD assays were strongly correlated (ELISA anti-RBD IgG and IgM/Pylon-IgG and IgM at 1.3 months, r=0.9200 and r=0.7543, p < 0.0001, respectively. Extended Data Fig 2a-d) while anti-N assays showed a moderate correlation (anti-N IgG ELISA and Roche anti-N total Ig at 1.3 months, r=0.3596, p = 0.0012. Extended Data Fig 2e-f). The anti-RBD and ELISA anti-N antibodies in plasma decreased significantly between 1.3 and 6.2 months (Fig. 1a-d). Notably, the decreased binding activity differed significantly by isotype and target. IgM showed the greatest decrease in anti-RBD reactivity (53%), followed by IgG (32%) while anti-RBD IgA decreased by only 15% and anti-N IgG levels by 22% (Fig. 1e). In contrast, the Roche anti-N assay^5^ showed a small but significant increase (19%) in reactivity between the two time points (Extended Data Fig 2g) which might be explained by the use of an antigen bridging approach^6^. In all cases the magnitude of the decrease was inversely proportional to and directly correlated with the initial antibody levels such that individuals with higher initial levels showed greater relative changes (Fig. 1f-i, Extended Data Fig 2h). All measurements were strongly correlated between the two time points (Extended Data Fig 2i-m) and anti-N IgG correlated with IgM, IgG and IgA anti-RBD reactivity at 1.3 months, respectively (Extended Data Fig. 2n-p). Notably, individuals with persistent post-acute symptoms had significantly higher anti-RBD IgG and anti-N total antibody levels at both study visits (Extended Data Fig. 1a-l).

**Fig. 1:**
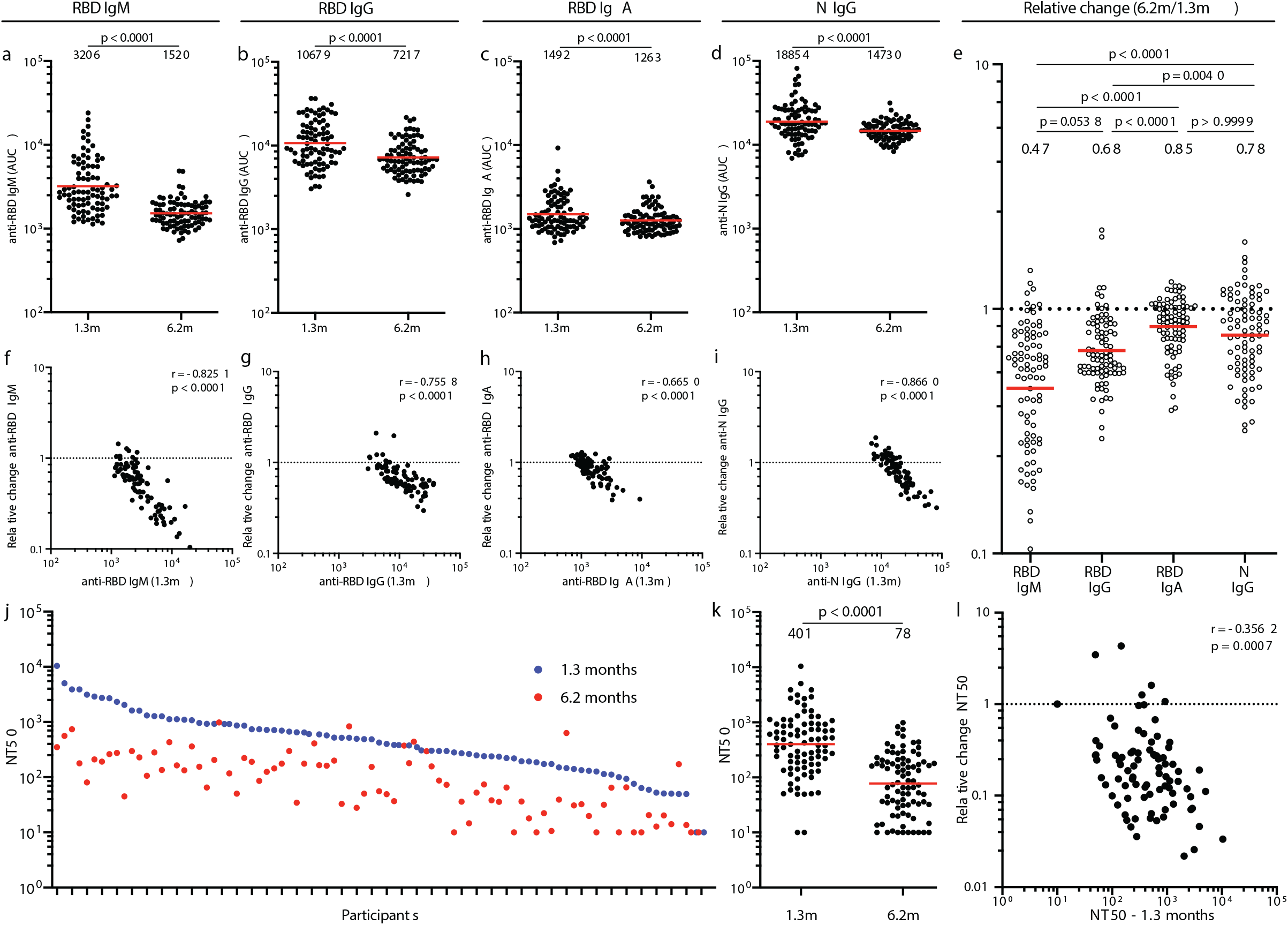
Plasma antibody dynamics against SARS-CoV-2. **a**–**d**, Results of ELISAs measuring plasma reactivity to RBD (**a**,**b,c**) and N protein (**d**) at the initial 1.3 and 6.2 month follow-up visit, respectively. **a**, Anti-RBD IgM. **b**, Anti-RBD IgG. **c**, Anti-RBD IgA **d**, Anti-N IgG. The normalized area under the curve (AUC) values for 87 individuals are shown in **a,b,c** and **d** for both time points, respectively. Positive and negative controls were included for validation ^1^. **e**, Relative change in plasma antibody levels between 1.3 and 6.2 months for anti-RBD IgM, IgG, IgA and anti-N IgG in all 87 individuals, respectively. **f-i,** Relative change in antibody levels between 1.3 and 6.2 months plotted against the corresponding antibody levels at 1.3 months. **f**, Anti-RBD IgM. r = −0.83, p <0.0001. **g**, Anti-RBD IgG. r = −0.76, p <0.0001. **h**, Anti-RBD IgA. r = −0.67, p <0.0001. **i**, Anti-N IgG. r = −0.87, p <0.0001. **j.** Ranked average half-maximal inhibitory plasma neutralizing titer (NT50) at 1.3 months (blue) and 6.2 months (red) for the 87 individuals studied. **k.** Graph shows NT50 for plasma from all 87 individuals collected at 1.3 and 6.2 months p <0.0001. **l.** Relative change in plasma neutralizing titers between 1.3 and 6.2 months plotted against the corresponding titers at 1.3 months. For **a**-**e**, **k** plotted values and horizontal bars indicate geometric mean. Statistical significance was determined using two-tailed Wilcoxon matched-pairs signed rank test in **a-d**, **k** and Friedman with Dunn’s multiple comparison test in **e**. The *r* and *p* values in **f** – **i** and **l** were determined by two-tailed Spearman’s correlations.

Plasma neutralizing activity was measured using an HIV-1 virus pseudotyped with the SARS-CoV-2 spike protein^1,7^. Consistent with other reports the geometric mean half-maximal neutralizing titer (NT_50_) in this group of 87 participants was 401 and 78 at 1.3 and 6.2 months, respectively, representing a five-fold decrease (Fig. 1j-k)^8,9^. Neutralizing activity was directly correlated with the IgG anti-RBD ELISA measurements (Extended data Fig. 2q-r). Moreover, the absolute magnitude of the decrease in neutralizing activity was inversely proportional to and directly correlated with the neutralizing activity at the earlier time point (Fig. 1l). We conclude that antibodies to RBD and plasma neutralizing activity decrease significantly but remain detectable 6 months after infection with SARS-CoV-2 in the majority of individuals.

To examine the phenotypic landscape of circulating B cells we performed high-dimensional flow cytometry on 41 randomly selected individuals at both time points and compared to pre-COVID-19 samples from healthy individuals (n=20). Global high-dimensional mapping with t-distributed stochastic neighbor embedding (tSNE) revealed significant persistent alterations in SARS-CoV-2 recovered individuals (Extended data Fig. 3a). The relative representation of clusters 2, 7, 8, 10 corresponding to naïve, memory, plasmablasts and plasma cells was decreased at 1.3 months and remained so at the later time point. Metacluster 15, which corresponds to immature B cells that are recent bone marrow immigrants was increased at the early time point but returned to control levels at the end of the observation period (Extended data Fig. 3b-d).

Whereas plasma cells are the source of circulating antibodies, memory B cells contribute to recall responses. To identify and enumerate the circulating SARS-CoV-2 memory B cell compartment we used flow cytometry to isolate individual B lymphocytes with receptors that bound to RBD ^1^ (Fig. 2a and Extended data Fig. 4). Notably, the percentage of RBD-binding memory B cells increased marginally between 1.3 and 6.2 months in 21 randomly selected individuals (Fig. 2b).

**Fig. 2:**
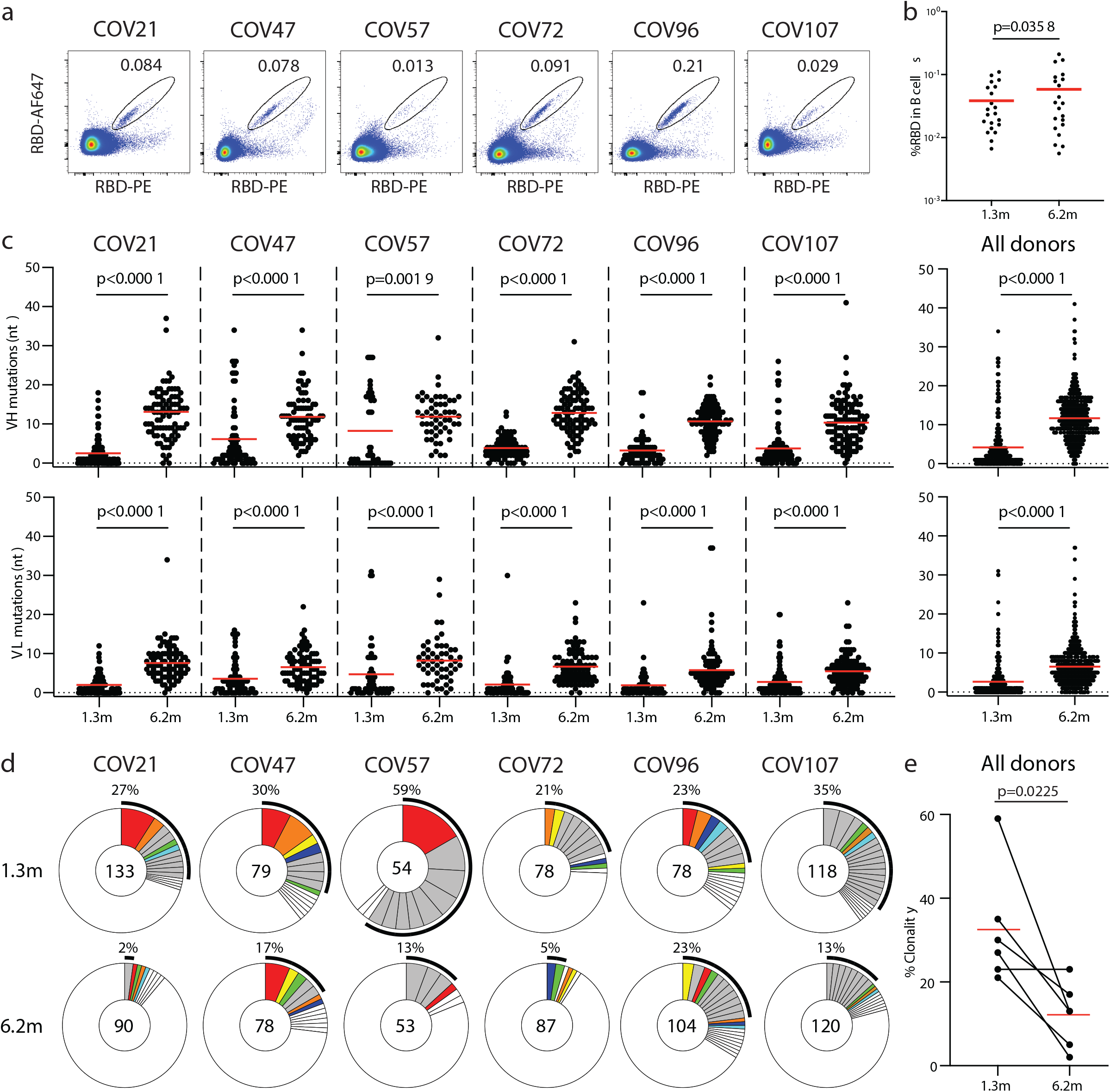
Anti-SARS-CoV-2 RBD antibody sequences. **a,** Representative flow cytometry plots showing dual AlexaFluor-647–RBD-and PE–RBD-binding B cells for six study individuals (gating strategy is in Extended Data Fig.5). Percentage of antigen-specific B cells is indicated. **b.** As in **a.** graph summarizes %RBD binding memory B cells in samples obtained at 1.3 and 6.2 months from 21 randomly selected individuals. Red horizontal bars indicate geometric mean values. Statistical significance was determined using two-tailed Wilcoxon matched-pairs signed rank test. **c,** Number of somatic nucleotide mutations in the IGVH (top) and IGVL (bottom) in antibodies obtained after 1.3 or 6.2 months from the indicated individual or all six donors (right). Statistical significance was determined using two-tailed Mann–Whitney U-tests and horizontal bars indicate median values. **d,** Pie charts show the distribution of antibody sequences from six individuals after 1.3 ^1^ (upper panel) or 6.2 months (lower panel). The number in the inner circle indicates the number of sequences analyzed for the individual denoted above the circle. Pie slice size is proportional to the number of clonally related sequences. The black outline indicates the frequency of clonally expanded sequences detected in each participant. Colored slices indicate persisting clones (same IGHV and IGLV genes and highly similar CDR3s) found at both timepoints in the same participant. Grey slices indicate clones unique to the timepoint. White slices indicate singlets found at both timepoints, while the remaining white area indicates sequences isolated once. **e.** Graph shows relative clonality at both time points for all six donors. Red horizontal bars indicate mean values. Statistical significance was determined using two-tailed t-test.

To determine whether there were changes in the antibodies produced by memory B cells after 6.2 months, we obtained 532 paired antibody heavy and light chains from the same 6 individuals that were examined at the earlier time point^1^ (Supplementary Table 3). There was no significant difference in IGV gene representation at the two time points, including the over-representation of the *IGHV3-30* and *3-53* gene segments ^1,10–15^ (Extended data Fig. 5a-b). In keeping with this observation, and similar to the earlier time point, antibodies that shared the same IGHV and IGLV genes comprised 8.6% of all sequences in different individuals (Extended data Fig. 5c). As might be expected, there was a small but significant overall increase in the percentage of IgG-expressing anti-RBD memory cells, from 49% to 58% (p=0.011, Extended data Fig. 5d-f). Consistent with the fractional increase in IgG memory cells, the extent of somatic hypermutation for both IGH and IGL differed significantly in all 6 individuals between the two time points. Whereas the average number of nucleotide mutations in IGH and IGL was only 4.2 and 2.8 at the first time point, these values were increased to 11.7 and 6.5 at the second time point (p<0.0001, Fig. 2c and Extended data Fig. 6a-f). In contrast, the overall average IGH and IGL CDR3 length and hydrophobicity were unchanged (Extended data Fig. 6g-h).

Similar to the earlier time point, we found expanded clones of memory B cells at 6.2 months including 23 that appeared at both time points. However, expanded clones accounted for only 12.4% of all antibody sequences after 6.2 months compared to 32% after 1.3 months (p = 0.0225, Fig. 2d-e). In addition, the overall clonal composition of the memory compartment differed at the two time points in all individuals examined (Fig. 2d). Forty-three expanded clones that were present at the earlier time point were not detectable after 6.2 months while 22 new expanded clones appeared. In addition, the relative distribution of clones that appeared at both time points also varied. For example, the dominant clones in COV21 and COV57 representing 9.0% and 16.7% of all sequences, respectively, were reduced to 1.1% and 1.9% of all sequences after 6.2 months (Fig. 2d and Supplementary Table 3). We conclude that while the magnitude of the RBD-specific memory B cell compartment is conserved between 1.3 and 6.2 months after SARS-CoV-2 infection, there is significant clonal turnover and antibody sequence evolution, consistent with prolonged germinal center reactions.

One hundred and twenty-two representative antibodies from the 6.2-month time point were tested for reactivity to RBD (Supplementary Table 4). The antibodies that were evaluated included: (1) 49 that were randomly selected from those that appeared only once; (2) 23 that appeared as singles at both 1.3 and 6.2 months; (3) 23 representatives of newly appearing expanded clones; (4) 27 representatives of expanded clones appearing at both time points. One hundred and fifteen of 122 of the antibodies bound to RBD indicating that flow cytometry efficiently identified B cells producing anti-RBD antibodies (Fig. 3a and Supplementary Tables 4 and 5). Taking all antibodies together, the mean ELISA half-maximal effective concentration (EC_50_) was not significantly different at the two time points (Fig. 3a, Supplementary Table 4 and 1). However, comparison of the antibodies that were present at both time points revealed a significant improvement of the EC_50_ after 6.2 months (p = 0.0227, Fig. 3b and Extended data Fig.7a).

**Fig. 3:**
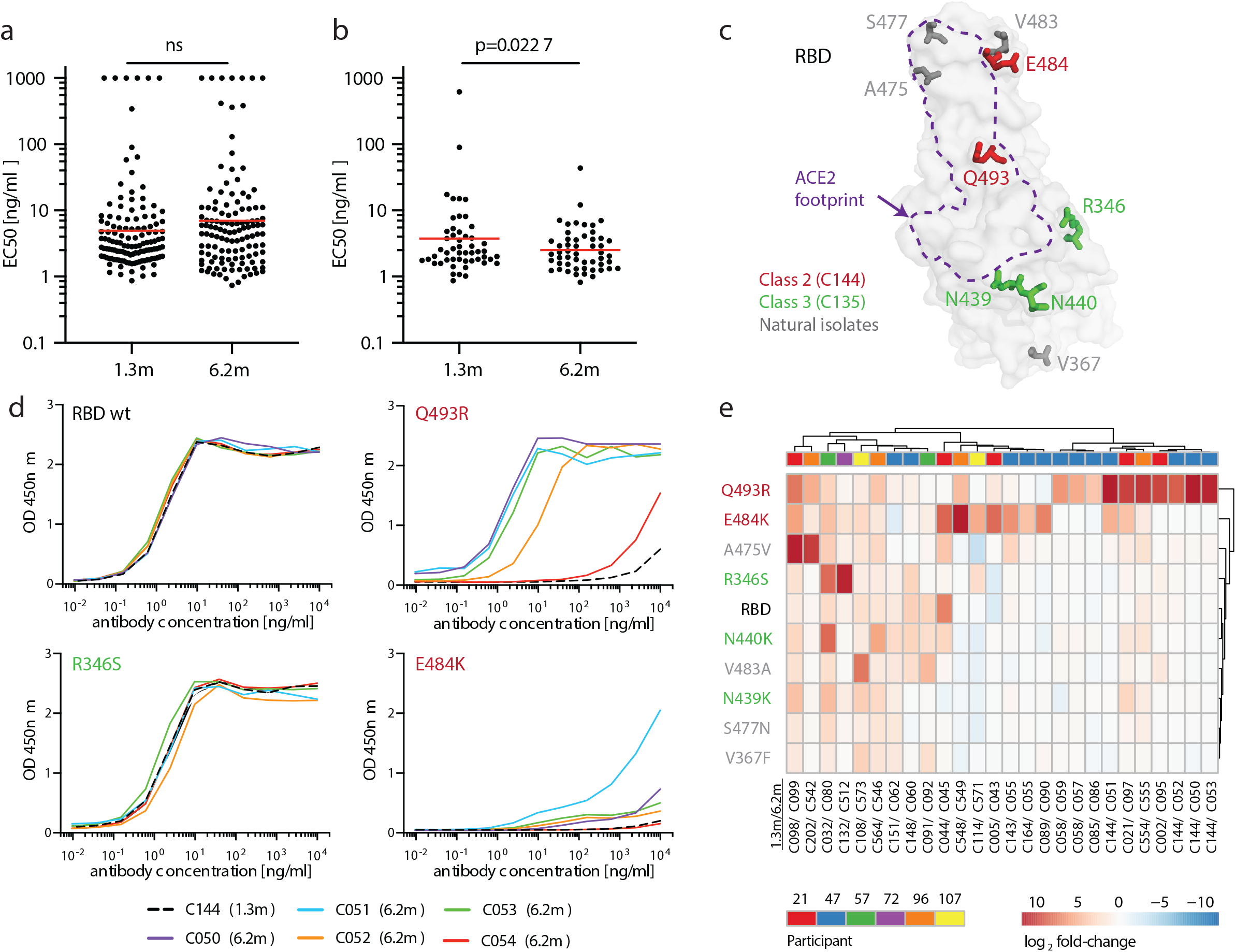
Anti-SARS-CoV-2 RBD monoclonal antibody reactivity. **a,** Graphs show anti-SARS-CoV-2 RBD antibody reactivity. ELISA EC_50_ values for all antibodies measured at 1.3 ^1^ and 122 selected monoclonal antibodies at 6.2 months. Horizontal bars indicate geometric mean. Statistical significance was determined using two-tailed Mann– Whitney U-test. **b**, EC_50_ values for all 52 antibodies that appear at 1.3 and 6.2 months. Average of two or more experiments. Horizontal bars indicate geometric mean. Statistical significance was determined using two-tailed Wilcoxon matched-pairs signed rank test. **c,** Surface representation of the RBD with the ACE2 binding footprint indicated as a dotted line and selected residues found in circulating strains (grey) and residues that mediate resistance to class 2 (red, C144) and 3 (green, C135) antibodies highlighted as sticks. **d.** Graphs show ELISA binding curves for C144 (black dashed line) and its clonal relatives obtained after 6.2 months (C050-54, solid lines) binding to wild type, Q493R, R346S, and E484K mutant RBDs. **e.** Heat map shows log2 relative fold change in EC_50_ against indicated RBD mutants for 26 antibody clonal pairs obtained at 1.3 and 6.2 month with the most pronounced changes in reactivity. The participant origin for each antibody pair is indicated above. All experiments were performed at least twice.

To determine whether the antibodies expressed by memory B cells at the late time point also showed altered breadth, we compared them to earlier clonal relatives in binding assays using control and mutant RBDs: The mutations E484K and Q493R^16^ were selected for resistance to class 2 antibodies such as C144 and C121 that bind directly to the ACE2 interaction ridge in the RBD ^1,17–19^ while R346S, N439K, and N440K were selected for resistance to class 3 antibodies such as C135 that do not directly interfere with ACE2 binding ^1,16–19^ (Fig. 3c). In addition, V367F, A475V, S477N, and V483A represent circulating variants that confer complete or partial resistance to class 1 and 2 antibodies ^16,17,20^ (Fig. 3c). Out of 52 antibody clonal pairs appearing at both time points, 43 (83%) showed overall increased binding to mutant RBDs at the 6.2-month time point (Extended data Fig. 7b-k, Supplementary Table 5). For example, C144, an antibody recovered at the 1.3-month time point, was unable to bind to Q493R or E484K RBDs, but all 5 of its 6.2-month clonal derivatives bound to Q493R, and one also showed binding to E484K (Fig. 3d). Overall, the most pronounced increase in binding occurred for RBD mutations in amino acid positions such as E484, Q493, N439, N440 and R346 that are critical for binding of class 2 and 3 antibodies ^16,17^ (Fig. 3e, Extended data Fig. 7b-k and Supplementary Table 5).

Next, all 122 antibodies from the 6.2 month time point were tested for activity in a pseudotyped SARS-CoV-2 neutralization assay ^1,7^ (Fig. 4a, Supplementary Table 6). Consistent with RBD binding assays, the mean neutralization half-maximal inhibitory concentrations (IC_50_) values were not significantly different at the two time points when all antibodies were compared (Fig. 4a and ^1^). However, comparison of the antibodies that were present at both time points revealed a significant improvement of the IC_50_ values at 6.2 months (p=0.0003, Fig. 4b and Extended data Fig. 8a).

**Fig. 4:**
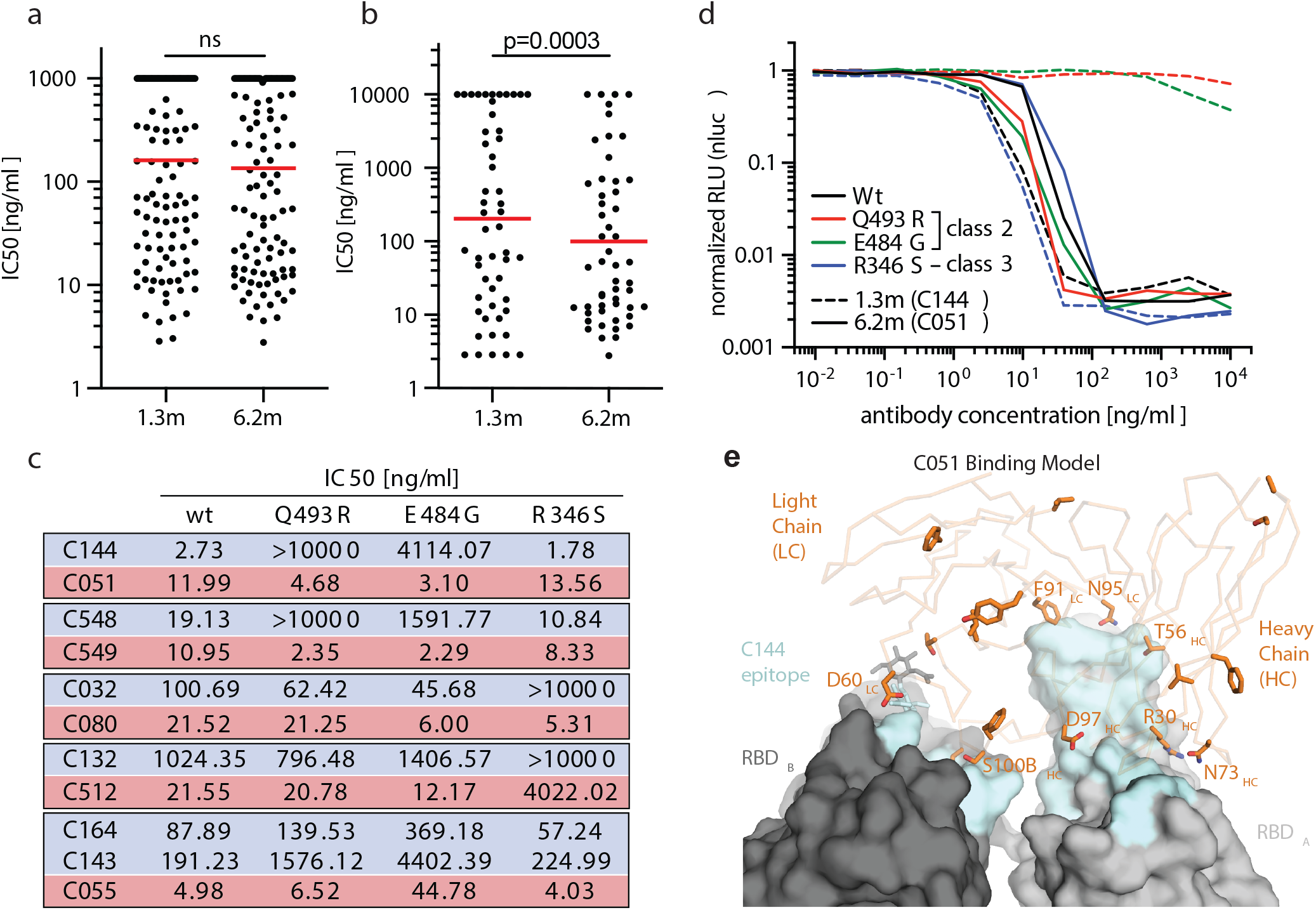
Anti-SARS-CoV-2 RBD monoclonal antibody neutralizing activity. **a,** SARS-CoV-2 pseudovirus neutralization assay. IC_50_ values for all antibodies measured at 1.3 months^1^ and 122 selected antibodies at 6.2 months. Antibodies with IC_50_ values above 1 µg/ml were plotted at 1 µg/ml. Mean of 2 independent experiments. Red bar indicates geometric mean. Statistical significance was determined using two-tailed Mann-Whitney U-test. **b,** IC50 values for 52 antibodies appearing at 1.3 and 6.2 months. Red bar indicates geometric mean. Statistical significance was determined using two-tailed Wilcoxon matched-pairs signed rank test. **c,** IC_50_ values for 5 different pairs of mAb clonal relatives obtained after 1.3 (blue) or 6.2 months (red) for neutralization of wild type and mutant SARS-CoV-2 pseudovirus. Antibody IDs of the 1.3 months/6.2 months mAb pairs as indicated. **d,** Graph shows the normalized relative luminescence values for cell lysates of 293T_ACE2_ cells 48 hpi with SARS-CoV-2 pseudovirus harboring wt RBD or three mutant RBDs (wt, Q493R, E484G, R346S RBD mutants are shown in black, red, green and blue, respectively) in the presence of increasing concentrations of two mAbs C144 (1.3 months, dashed lines) or C051 (6.2 months, solid lines). Experiments were performed at least twice. **e,** Surface representation of two adjacent “down” RBDs (RBD_A_ and RBD_B_) on a spike trimer with the C144 epitope on the RBDs highlighted in cyan and positions of amino acid mutations that accumulated in C051 compared to the parent antibody C144 highlighted as stick side chains on a Cα atom representation C051 V_H_V_L_ binding to adjacent RBDs. The C051 interaction with two RBDs was modeled based on a cryo-EM structure of C144 Fab bound to spike trimer ^17^.

To determine whether the antibodies exhibiting altered RBD binding also show increased neutralizing breadth, we tested 5 representative antibody pairs recovered at the two time points against HIV-1 viruses pseudotyped with E484G, Q493R, and R346S mutant spike proteins (Fig. 4c, Supplementary Table 6). Notably, the Q493R and E484G pseudotyped viruses were resistant to neutralization by C144; in contrast, its 6.2-month clonal derivative C051 neutralized both variants with IC_50_ values of 4.7 and 3.1 ng/ml respectively (Fig. 4c-d). Similarly, R346S pseudotyped viruses were resistant to C032, but a 6.2-month clonal derivative C080 neutralized this variant with an IC_50_ of 5.3 ng/ml (Fig. 4c, Extended data Fig. 8b-f). Consistent with the observed changes in binding and neutralizing activity several late-appearing antibodies (e.g. C051) had acquired mutations directly in or adjacent to the RBD-binding paratope (Fig. 4e, Extended data Fig. 8g-j). We conclude that memory B cells that evolved during the observation period express antibodies with increased neutralizing potency and breadth.

Antibody evolution occurs by somatic mutation and selection in germinal centers wherein antigen can be retained in the form of immune complexes on the surface of follicular dendritic cells for prolonged periods of time. Residual protein in tissues represents another potential source of antigen. SARS-CoV-2 replicates in ACE2-expressing cells in the lungs, nasopharynx and small intestine ^21–24^, and viral RNA has been detected in stool samples even after the virus is cleared from the nasopharynx ^25–27^. To determine whether there might be antigen persistence in the intestine after resolution of clinical illness, we obtained biopsies from the upper and lower gastrointestinal (GI) tract of 14 individuals, an average of 4 months (range 2.8-5.7 months) after initial SARS-CoV-2 diagnosis (Supplementary Table 7). Immunostaining was performed to determine whether viral protein was also detectable in upper and lower GI tract, with de-identified biopsies from individuals pre-dating the pandemic (n=10) serving as controls. ACE2 and SARS-CoV-2 N protein was detected in intestinal enterocytes in 5 of 14 individuals (Fig. 5a-d, Extended data Fig. 9a-h and 10a-b, and Supplementary Table 7) but not in historic control samples (Extended data Fig. 9i-l). When detected, immunostaining was sporadic, patchy, exclusive to the intestinal epithelium and not associated with inflammatory infiltrates (Extended data Fig. 9a-h and 10a-b). Clinically approved nasopharyngeal swab PCR assays were negative in all 14 individuals at the time of biopsy. However, biopsy samples from 3 of the 14 participants produced PCR amplicons that were sequence verified as SARS-CoV-2 (Methods and Supplementary Table 7). In addition, viral RNA was detected by in situ hybridization in biopsy samples from the two participants that were tested (Extended data Fig. 10c-d) but not in historic control samples (Extended data Fig. 10e). Sampling variability during the endoscopic procedure likely contributed to incomplete concordance between detection of viral RNA and protein assays.

**Fig. 5:**
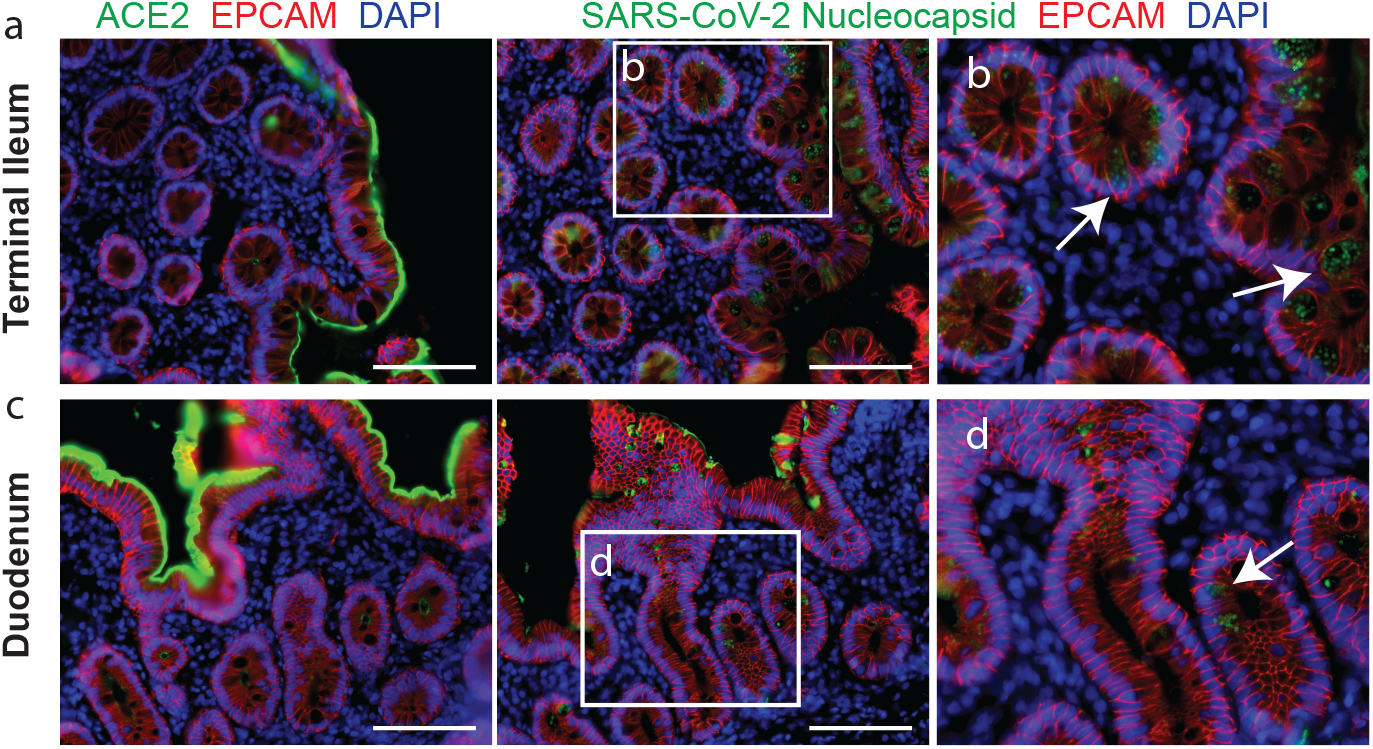
Immunofluorescence imaging of intestinal biopsies. **a,** Immunofluorescence images of human enterocytes stained for EPCAM (red), DAPI (blue) and either ACE2 (green, **a** and **c**) or SARS-CoV-2 N (green, **b** and **d**) in intestinal biopsies taken 92 days after COVID-19 symptom onset of participant CGI-088 in the terminal ileum (**a-b**) or duodenum (**c-d**). Arrows indicate enterocytes with detectable SARS-CoV-2 antigen. White scale bar corresponds to 100 μm. The experiments were repeated independently at least twice with similar results.

Neutralizing antibodies to SARS-CoV-2 develop in most individuals after infection but decay with time ^6,8,9,28–31^. These antibodies are effective in prevention and therapy in animal models and are likely to play a role in protection from re-infection in humans ^32^. Although there is a significant decrease in plasma neutralizing activity between 1.3 and 6.2 months, antibody titers remain measurable in most individuals ^8,9,28–31,33–36^.

Neutralizing monoclonal antibodies obtained from individuals during the early convalescence period showed remarkably low levels of somatic mutations that some investigators attributed to defects in germinal center formation ^10,11,13,37–40^. Our data indicate that the anti-SARS-CoV-2 memory B cell response evolves during the first 6 months after infection, with accumulation of Ig somatic mutations, and production of antibodies with increased neutralizing breadth and potency. Persistent antibody evolution occurs in germinal centers and requires that B cells are exposed to antigen trapped in the form of immune complexes on follicular dendritic cells ^41^. This form of antigen can be long-lived because follicular dendritic cells do not internalize immune complexes. In addition, even small amounts of persistent viral antigen could fuel antibody evolution. The observation that SARS-CoV-2 mRNA and protein remains detectable in the small intestinal epithelium in some individuals months after infection is consistent with the relative persistence of anti-RBD IgA antibodies and continued antibody evolution ^34–36^.

Memory responses are responsible for protection from re-infection and are essential for effective vaccination. The observation that memory B cell responses do not decay after 6.2 months ^34,35,42^, but instead continue to evolve, is strongly suggestive that individuals who are infected with SARS-CoV-2 could mount a rapid and effective response to the virus upon re-exposure.

## Methods

### Data reporting

No statistical methods were used to predetermine sample size. The experiments were not randomized and the investigators were not blinded to allocation during experiments and outcome assessment.

#### Study participants

Previously enrolled study participants ^1^ were asked to return for a 6-month follow-up visit at the Rockefeller University Hospital in New York from August 31 through October 16, 2020. Eligible participants were adults aged 18-76 years and were either diagnosed with SARS-CoV-2 infection by RT-PCR (cases), or were close contacts (e.g., household, co-workers, members of same religious community) with someone who had been diagnosed with SARS-CoV-2 infection by RT-PCR (contacts). Close contacts without seroconversion against SARS-CoV-2 as assessed by serological assays (described below) were not included in the subsequent analysis. Most study participants were residents of the Greater New York City tri-state region and were asked to return approximately 6 months after the time of onset of COVID-19 symptoms. Participants presented to the Rockefeller University Hospital for blood sample collection and were asked to recall the symptoms and severity of clinical presentation during the acute (first 6 weeks) and the convalescent (7 weeks until second study visit) phase of COVID-19, respectively. The severity of acute infection was assessed by the WHO Ordinal Clinical Progression/Improvement Scale (https://www.who.int/publications/i/item/covid-19-therapeutic-trial-synopsis). Shortness of breath was assessed through the modified Medical Research Council (mMRC) dyspnea scale ^43^. Participants who presented with persistent symptoms attributable to COVID-19 were identified on the basis of chronic shortness of breath or fatigue, deficit in athletic ability and/or three or more additional long-term symptoms such as persistent unexplained fevers, chest pain, new-onset cardiac sequalae, arthralgias, impairment of concentration/mental acuity, impairment of sense of smell/taste, neuropathy or cutaneous findings ^2,3^. All participants at Rockefeller University provided written informed consent before participation in the study and the study was conducted in accordance with Good Clinical Practice. Clinical data collection and management were carried out using the software iRIS by iMedRIS. The study was performed in compliance with all relevant ethical regulations and the protocol for studies with human participants was approved by the Institutional Review Board of the Rockefeller University.

#### Gastrointestinal biopsy cohort

To determine if SARS-CoV-2 can persist in the gastrointestinal tract, we recruited a cohort of 14 individuals with prior diagnosis of and recovery from COVID-19 illness. Eligible participants included adults, 18-76 years of age who were previously diagnosed with SARS-CoV-2 by RT-PCR or through a combination of clinical symptoms consistent with COVID-19 plus evidence of seroconversion, and presented to the gastroenterology clinics of Mount Sinai Hospital. Endoscopic procedures were performed for clinically indicated conditions as detailed in Supplementary Table 7. All participants were asymptomatic at the time of the endoscopic procedures and negative for SARS-CoV-2 by nasal swab PCR (Cycle threshold (Ct) cut-off <38).

The CLIA certified laboratory of the Mount Sinai Health System validated the laboratory-developed nasopharyngeal swab real time RT-PCR test according to the New York State Department of Health Wadsworth Center validation procedure for SARS-CoV-2^44^. Informed consent was obtained from all participants. The biopsy-related studies were approved by the Mount Sinai Ethics Committee/IRB (IRB 16-0583, The impact of viral infections and their treatment on gastrointestinal immune cells).

### SARS-CoV-2 saliva PCR test

The SARS-CoV-2 PCR method for saliva samples was developed and its performance characteristics determined by the Rockefeller University Clinical Genomics Laboratory. This laboratory-developed test has been authorized by New York State under an Emergency Use Authorization (EUA) for use by authorized laboratories. Saliva was collected into guanidine thiocyanate buffer as described ^45^. RNA was extracted using either a column-based (Qiagen QIAmp DSP Viral RNA Mini Kit, Cat#61904) or a magnetic bead-based method as described ^46^. Reverse transcribed cDNA was amplified using primers and probes validated by the CDC or by Columbia University Personalized Medicine Genomics Laboratory, respectively, and approved by the FDA under the Emergency Use Authorization. Viral RNA was considered detected if the cycle threshold (Ct) for two viral primers/probes were <40.

#### Blood samples processing and storage

Peripheral Blood Mononuclear Cells (PBMCs) were obtained by gradient centrifugation and stored in liquid nitrogen in the presence of FCS and DMSO. Heparinized plasma and serum samples were aliquoted and stored at −20°C or less. Prior to experiments, aliquots of plasma samples were heat-inactivated (56°C for 1 hour) and then stored at 4°C.

### High throughput automated serology assays

Plasma samples from 80 out of 87 participants were tested by high throughput automated serology assays. The Roche Elecsys anti-SARS-CoV-2 assay was performed on Roche Cobas e411 (Roche Diagnostics, Indianapolis, IN). The Elecsys antinSARSnCoV-2 assay uses a recombinant protein representing the N antigen for the determination of antibodies against SARSnCoVn2. This assay received Emergency Use Authorization (EUA) approval from the United States Food and Drug Administration (FDA) ^5^. The Pylon COVID-19 IgG and IgM assays were used to measure plasma IgG and IgM antibodies against SARS-CoV-2, respectively. Plasma samples were assayed on the Pylon 3D analyzer (ET HealthCare, Palo Alto, CA) as previously described ^4^. This assay was implemented clinically as a laboratory-developed test under New York State Department of Health regulations. Briefly, the assay was performed using a unitized test strip containing wells with pre-dispensed reagents. The COVID-19 reagent contains biotinylated recombinant versions of the SARS-CoV-2 S-Protein RBD and trace amounts of N protein as antigens that bind IgG and IgM, respectively. The cut off values for both Pylon assays were determined using the mean of non-COVID-19 samples plus 6 Standard Deviations (SDs). The results of a sample are reported in the form of a cutoff index (COI) or an index value (IV), which were determined by the instrument readout of the test sample divided by instrument readout at cut off.

### ELISAs

ELISAs ^47,48^ to evaluate antibodies binding to SARS-CoV-2 N (Sino Biological 40588-V08B), RBD and additional RBDs were performed by coating of high-binding 96-half-well plates (Corning 3690) with 50 μl per well of a 1μg/ml protein solution in PBS overnight at 4 °C. Plates were washed 6 times with washing buffer (1× PBS with 0.05% Tween-20 (Sigma-Aldrich)) and incubated with 170 μl per well blocking buffer (1× PBS with 2% BSA and 0.05% Tween-20 (Sigma)) for 1 h at room temperature. Immediately after blocking, monoclonal antibodies or plasma samples were added in PBS and incubated for 1 h at room temperature. Plasma samples were assayed at a 1:67 starting dilution and 7 additional threefold serial dilutions. Monoclonal antibodies were tested at 10 μg/ml starting concentration and 10 additional fourfold serial dilutions. Plates were washed 6 times with washing buffer and then incubated with anti-human IgG, IgM or IgA secondary antibody conjugated to horseradish peroxidase (HRP) (Jackson Immuno Research 109-036-088 109-035-129 and Sigma A0295) in blocking buffer at a 1:5,000 dilution (IgM and IgG) or 1:3,000 dilution (IgA). Plates were developed by addition of the HRP substrate, TMB (ThermoFisher) for 10 min (plasma samples) or 4 minutes (monoclonal antibodies), then the developing reaction was stopped by adding 50 μl 1 M H_2_SO_4_ and absorbance was measured at 450 nm with an ELISA microplate reader (FluoStar Omega, BMG Labtech) with Omega and Omega MARS software for analysis. For plasma samples, a positive control (plasma from participant COV72, diluted 66.6-fold and seven additional threefold serial dilutions in PBS) was added to every assay plate for validation. The average of its signal was used for normalization of all of the other values on the same plate with Excel software before calculating the area under the curve using Prism V8.4 (GraphPad). For monoclonal antibodies, the EC50 was determined using four-parameter nonlinear regression (GraphPad Prism V8.4).

### Expression of RBD proteins

Mammalian expression vectors encoding the RBDs of SARS-CoV-2 (GenBank MN985325.1; S protein residues 319-539) and eight additional mutant RBD proteins (E484K, Q493R, R346S, N493K, N440K, V367F, A475V, S477N and V483A) with an N-terminal human IL-2 or Mu phosphatase signal peptide were previously described ^49^.

### SARS-CoV-2 pseudotyped reporter virus

SARS-CoV-2 pseudotyped particles were generated as previously described ^1,50^. Briefly, 293T cells were transfected with pNL4-3ΔEnv-nanoluc and pSARS-CoV-2-S_Δ19_. For generation of RBD-mutant pseudoviruses, pSARS-CoV-2-S _Δ19_ carrying either of the following spike mutations was used instead of its wt counterpart: Q493R, R346S or E484G ^51^. Particles were harvested 48 hpt, filtered and stored at −80°C.

### Pseudotyped virus neutralization assay

Fourfold serially diluted plasma from COVID-19-convalescent individuals or monoclonal antibodies were incubated with SARS-CoV-2 pseudotyped virus for 1 h at 37 °C. The mixture was subsequently incubated with 293T_Ace2_ cells for 48 h after which cells were washed with PBS and lysed with Luciferase Cell Culture Lysis 5× reagent (Promega). Nanoluc Luciferase activity in lysates was measured using the Nano-Glo Luciferase Assay System (Promega) with the Glomax Navigator (Promega). The obtained relative luminescence units were normalized to those derived from cells infected with SARS-CoV-2 pseudotyped virus in the absence of plasma or monoclonal antibodies. The half-maximal inhibitory concentration for plasma (NT_50_) or monoclonal antibodies (IC_50_) was determined using four-parameter nonlinear regression (least squares regression method without weighting; constraints: top=1, bottom=0) (GraphPad Prism).

### High-dimensional data analysis of flow cytometry data

viSNE and FlowSOM analyses were performed on B cells using the Cytobank platform (https://cytobank.org). viSNE analysis was performed using equal sampling of 4893 cells from each FCS file, with 7500 iterations, a perplexity of 30, and a theta of 0.5. The following markers were used to generate viSNE maps: IgA, CD305, TGFb-RII, CD138, CD10, CD272, IgD, CD24, CD21, CD95, HLA-DR, IgG, CD279, CD38, IgM, CD274, CD27, CD23, CXCR5, CD32, CD86, CD40, CD85j, CD11c, and CXCR3. Resulting viSNE maps were fed into the FlowSOM clustering algorithm^52^. The self-organizing map (SOM) was generated using hierarchical consensus clustering on the tSNE axes.

### Heatmap visualization

Heatmaps to display column-scaled z-scores of MFI for individual FlowSOM clusters according to marker expression were created using the R function pheatmap.

### Biotinylation of viral protein for use in flow cytometry

Purified and Avi-tagged SARS-CoV-2 RBD was biotinylated using the Biotin-Protein Ligase-BIRA kit according to manufacturer’s instructions (Avidity) as described before ^1^. Ovalbumin (Sigma, A5503-1G) was biotinylated using the EZ-Link Sulfo-NHS-LC-Biotinylation kit according to the manufacturer’s instructions (Thermo Scientific). Biotinylated ovalbumin was conjugated to streptavidin-BV711 (BD biosciences, 563262) and RBD to streptavidin-PE (BD Biosciences, 554061) and streptavidin-AF647 (Biolegend, 405237)^1^.

### Single-cell sorting by flow cytometry

Single-cell sorting by flow cytometry was described previously ^1^. Briefly, peripheral blood mononuclear cells were enriched for B cells by negative selection using a pan-B-cell isolation kit according to the manufacturer’s instructions (Miltenyi Biotec, 130-101-638). The enriched B cells were incubated in FACS buffer (1× PBS, 2% FCS, 1 mM EDTA) with the following anti-human antibodies (all at 1:200 dilution): anti-CD20-PECy7 (BD Biosciences, 335793), anti-CD3-APC-eFluro 780 (Invitrogen, 47-0037-41), anti-CD8-APC-eFluor 780 (Invitrogen, 47-0086-42), anti-CD16-APC-eFluor 780 (Invitrogen, 47-0168-41), anti-CD14-APC-eFluor 780 (Invitrogen, 47-0149-42), as well as Zombie NIR (BioLegend, 423105) and fluorophore-labelled RBD and ovalbumin (Ova) for 30 min on ice. Single CD3−CD8−CD14−CD16−CD20+Ova−RBD-PE+RBD-AF647+ B cells were sorted into individual wells of 96-well plates containing 4 μl of lysis buffer (0.5× PBS, 10 mM DTT, 3,000 units/ml RNasin Ribonuclease Inhibitors (Promega, N2615) per well using a FACS Aria III and FACSDiva software (Becton Dickinson) for acquisition and FlowJo for analysis. The sorted cells were frozen on dry ice, and then stored at −80 °C or immediately used for subsequent RNA reverse transcription.

### Antibody sequencing, cloning and expression

Antibodies were identified and sequenced as described previously ^1^. In brief, RNA from single cells was reverse-transcribed (SuperScript III Reverse Transcriptase, Invitrogen, 18080-044) and the cDNA stored at −20 °C or used for subsequent amplification of the variable IGH, IGL and IGK genes by nested PCR and Sanger sequencing. Sequence analysis was performed using MacVector. Amplicons from the first PCR reaction were used as templates for sequence-and ligation-independent cloning into antibody expression vectors. Recombinant monoclonal antibodies and Fabs were produced and purified as previously described ^1^.

### Computational analyses of antibody sequences

Antibody sequences were trimmed based on quality and annotated using Igblastn v.1.14. with IMGT domain delineation system. Annotation was performed systematically using Change-O toolkit v.0.4.540 ^53^. Heavy and light chains derived from the same cell were paired, and clonotypes were assigned based on their V and J genes using in-house R and Perl scripts (Extended data Fig. 5). All scripts and the data used to process antibody sequences are publicly available on GitHub (https://github.com/stratust/igpipeline).

The frequency distributions of human V genes in anti-SARS-CoV-2 antibodies from this study was compared to 131,284,220 IgH and IgL sequences generated by ^54^ and downloaded from cAb-Rep^55^, a database of human shared BCR clonotypes available at https://cab-rep.c2b2.columbia.edu/. Based on the 82 distinct V genes that make up the 1703 analyzed sequences from Ig repertoire of the three participants present in this study, we selected the IgH and IgL sequences from the database that are partially coded by the same V genes and counted them according to the constant region. The frequencies shown in (Fig. S4) are relative to the source and isotype analyzed. We used the two-sided binomial test to check whether the number of sequences belonging to a specific IgHV or IgLV gene in the repertoire is different according to the frequency of the same IgV gene in the database. Adjusted p-values were calculated using the false discovery rate (FDR) correction. Significant differences are denoted with stars.

Nucleotide somatic hypermutation and CDR3 length were determined using in-house R and Perl scripts. For somatic hypermutations, IGHV and IGLV nucleotide sequences were aligned against their closest germlines using Igblastn and the number of differences were considered nucleotide mutations. The average mutations for V genes was calculated by dividing the sum of all nucleotide mutations across all participants by the number of sequences used for the analysis. To calculate the GRAVY scores of hydrophobicity ^56^ we used Guy H.R. Hydrophobicity scale based on free energy of transfer (kcal/mole) ^57^ implemented by the R package Peptides (the Comprehensive R Archive Network repository; https://journal.r-project.org/archive/2015/RJ-2015-001/RJ-2015-001.pdf). We used 532 heavy chain CDR3 amino acid sequences from this study and 22,654,256 IGH CDR3 sequences from the public database of memory B cell receptor sequences ^58^. The Shapiro–Wilk test was used to determine whether the GRAVY scores are normally distributed. The GRAVY scores from all 532 IGH CDR3 amino acid sequences from this study were used to perform the test and 5,000 GRAVY scores of the sequences from the public database were randomly selected. The Shapiro–Wilk P values were 6.896 × 10−3 and 2.217 × 10−6 for sequences from this study and the public database, respectively, indicating that the data were not normally distributed. Therefore, we used the two-tailed Wilcoxon nonparametric test to compare the samples, which indicated a difference in hydrophobicity distribution (P = 5 × 10−6) (Extended data Fig. 6h).

Heatmap of log2 relative fold change in EC50 against the indicated RBD mutants for antibody clonal pairs obtained at 1.3 and 6.2 months (Fig.3e and Extended data Fig. 7k) was created with R pheatmap package (https://github.com/raivokolde/pheatmap) using Euclidean distance and Ward.2 clustering method.

### Biopsies and Immunofluorescence

Endoscopically obtained mucosal biopsies were formalin fixed and paraffin embedded. Sections (5µm) were cut, dewaxed in xylene, and rehydrated in graded alcohol and phosphate-buffered saline (PBS). Heat-induced epitope retrieval was performed in target retrieval solution (Dako, S1699) using a commercial pressure cooker. Slides were then cooled to room temperature, washed in PBS and permeabilized for 30 minutes in 0.1% Triton X-100 in PBS. Non-specific binding was blocked with 10% goat serum (Invitrogen, 50062Z) for 1 hour at room temperature. Sections were then incubated with a combination of primary antibodies diluted in blocking solution overnight at 4°C. Slides were washed 3 times in PBS and then incubated in secondary antibody and 4′,6-diamidino-2-phenylindole (1ug/mL) for 1 hour at room temperature. Sections were washed in PBS 3 times and then mounted with Fluoromount-G (Electron Microscopy Sciences, 1798425). Controls included, omitting primary antibody (no primary 995 control), or substituting primary antibodies with non-reactive antibodies of the same isotype (isotype control). A Nikon Eclipse Ni microscope and digital SLR camera (Nikon, DS-Qi2) was used to visualize and image the tissue.

The antibody used to stain sections for N protein was raised in rabbits against SARS-CoV N and is cross-reactive with SARS-CoV-2 N protein ^59^.

| Antigen | Clone | Vendor | Catalogue number | Host species | Conjugate | Dilution |
| --- | --- | --- | --- | --- | --- | --- |
| ACE2 | Polyclonal | Abcam | ab15348 | rabbit | unconjugated | 1:1000 |
| EPCAM | SPM491 | GeneTex | GTX34693 | mouse | unconjugated | 1:100 |
| SARS-CoV-2 nucleocapsid | Polyclonal | Spiegel, M. *et al.*<sup>59</sup> | N/A | rabbit | unconjugated | 1:2000 |
| No known specificity (isotype control) | Polyclonal | Abcam | ab37415 | rabbit | unconjugated | 1:100 to 1:2000 corresponding to the concentration of the targeted polyclonal rabbit antibody |
| Yeast GAL4 (isotype control) | 15-6E10A7 | Abcam | ab170190 | mouse | unconjugated | 1:100 to 1:2000 corresponding to the concentration of the targeted monoclonal mouse antibody |
| Mouse IgG H&L | Polyclonal | Abcam | ab150116 | goat | Alexa Fluor 594 | 1:1000 |
| Rabbit IgG H&L | Polyclonal | Abcam | ab150077 | goat | Alexa Fluor 488 | 1:1000 |

### SARS-CoV-2 PCR (intestinal biopsies)

To determine if SARS-CoV-2 RNA is present in the gastrointestinal tract we isolated RNA from endoscopically obtained mucosal biopsies using Direct-zol miniprep kit (Zymo research, Cat. No. R2050). Reverse transcribed cDNA was amplified using 2019-nCov Ruo Kit (IDT) to detect viral nucleocapsid genomic RNA. Amplification of sub-genomic nucleocapsid RNA was done using following primers and probe: sgLeadSARSCov2_F 5’-CGATCTCTTGTAGATCTGTTCTC -3’^27^, wtN_R4 5’ – GGTGAACCAAGACGCAGTAT – 3’, wtN_P4 5’ – /56-FAM/TAACCAGAA/ZEN/TGGAGAACGCAGTGGG/3IABkFQ/ – 3’. Quantitative PCR was performed using QuantTect probe PCR kit (Qiagen, Cat. No. 204345) under following conditions: 95 15’, 95°C 15 sec, 60°C 1 min using the Applied Biosystem QuantStudio 6 Flex Real-Time PCR System. Viral RNA was considered detected if the cycle threshold (Ct) for viral primers/probes was <40. Samples from positive wells were column purified and presence of N1 sequences additionally verified by Sanger sequencing.

### SARS-CoV-2 RNA detection by probes proximity ligation

Probes were designed with a 20-25 nucleotides homology to SARS-CoV-2 genomic RNA. Probes were assessed by NCBI BLAST to exclude off target binding to other cellular transcripts. IDT OligoAnalyzer (Integrated DNA Technologies) was used to identify probe pairs with similar thermodynamic properties; melting temperature 45-60°C, GC content of 40-55%, and low self-complementary. The 3’ end of each one of the probes used for proximity ligation signal amplification is designed with a partially complementary sequence to the 61bp long backbone and partially to the 21bp insert as previously described (Supplementary Table 8). smFISH probes were designed with a complementary 3’ end to the biotin detection probe (Supplementary Table 8).

Paraffin embedded samples were sectioned at ten microns. Sections were deparaffinized using 100% xylene, 5 min at room temperature, repeated twice. Slides were rinsed in 100% ethanol, 1 min at room temperature, twice and air dried. Endogenous peroxidase activity was eliminated by treating the samples with 0.3% hydrogen peroxide, 10 min at room temperature followed by washing with DEPC treated water. Samples were incubated 15 min at 95-100 °C in antigen retrieval solution (ACDBio, Newark, CA, USA) rinsed in DEPC treated water and dehydrated in 100% ethanol, 3 min at room temperature and air dried. Tissue sections were permeabilized 30 min at 40°C using RNAscope protease plus solution (ACDBio, Newark, CA, USA) and rinsed in DEPC treated water.

Hybridization was performed overnight at 40°C in a buffer based on DEPC-treated water containing 2× SSC, 20% formamide (Thermo Fischer Scientific, Waltham, MA, USA), 2.5 % (vol/vol) polyvinylsulfonic acid, 20 mM ribonucleoside vanadyl complex (New England Biolabs, Ipswich, MA, USA), 40 U/ml RNasin (Promega, Madison, WI, USA), 0.1% (vol/vol) Tween 20 (Sigma Aldrich), 100 μg/ml salmon sperm DNA (Thermo Fisher Scientific, Waltham, MA, USA), 100 μg/ml yeast RNA (Thermo Fisher Scientific, Waltham, MA, USA). DNA probes dissolved in DEPC-treated water were added at a final concentration of 100nM (Integrated DNA Technologies, Coralville, IA, ISA). Samples were washed briefly and incubated in a buffer containing, 2× SSC, 20% formamide, 40 U/ml RNasin at 40 °C and then washed four times (5 min each) in wash buffer, PBS, 0.1% (vol/vol) Tween 20, and 4 U/ml RNasin (Promega, Madison, WI, USA). Slides were then incubated with 100 nM insert/backbone oligonucleotides in PBS, 1× SSC, 0.1% (vol/vol) Tween 20, 100 μg/ml salmon sperm DNA (Thermo Fisher Scientific, Waltham, MA, USA), 100 μg/ml yeast RNA (Thermo Fisher Scientific, Waltham, MA, USA), 40 U/ml RNasin at 37 °C. After four washes, tissues were incubated at 37°C with 0.1 U/µl T4 DNA ligase (New England Biolabs, Ipswich, MA, USA) in 50mM Tris-HCl, 10mM MgCl2, 1mM ATP, 1mM DTT, 250µg/ml BSA, 0.05% Tween 20, 40 U/ml RNasin, followed by incubation with 0.1 U/µl phi29 DNA polymerase in 50 mM Tris–HCl, 10 mM MgCl2, 10 mM (NH4)2SO4, 250nM dNTPs, 1mM DTT, 0.05% Tween 20, 40 U/ml RNasin pH 7.5 at 30 °C. Slides were washed and endogenous biotin was blocked using Avidin/Biotin blocking kit (Vector laboratories, Burlingame, CA, USA) according to the manufacture instructions. Rolling cycle amplicons were identified using a biotin labeled DNA probe at a concentration of 5 nM at 37 °C in PBS, 1× SSC, 0.1% Tween 20, 100 μg/ml salmon sperm DNA, 100 μg/ml yeast RNA. After washing, samples were incubated with 1:100 diluted streptavidin-HRP (Thermo Fisher Scientific, Waltham, MA, USA) in PBS, 60 min at room temperature followed by washing. Fluorescent labeling was accomplished using Alexa Fluor 647 Tyramide SuperBoostKit (Thermo Fischer Scientific, Waltham, MA, USA) according to manufacture instructions. Hoechst 33342 was used for nuclear counterstaining (Thermo Fischer Scientific, Waltham, MA, USA) and samples were mounted in ProLong gold antifade (Thermo Fischer Scientific, Waltham, MA, USA).

### SARS-CoV-2 RNA detection by smFISH

Hybridization was performed overnight at 40°C in a buffer based on DEPC-treated water containing 2× SSC, 20% formamide (Thermo Fischer Scientific, Waltham, MA, USA), 2.5 % (vol/vol) polyvinylsulfonic acid, 20 mM ribonucleoside vanadyl complex (New England Biolabs, Ipswich, MA, USA), 40 U/ml RNasin (Promega, Madison, WI, USA), 0.1% (vol/vol) Tween 20 (Sigma Aldrich), 100 μg/ml salmon sperm DNA (Thermo Fisher Scientific, Waltham, MA, USA), 100 μg/ml yeast RNA (Thermo Fisher Scientific, Waltham, MA, USA). DNA probes dissolved in DEPC-treated water were added at a final concentration of 10nM (Integrated DNA Technologies, Coralville, IA, ISA). Samples were washed briefly and incubated in a buffer containing, 2× SSC, 20% formamide, 40 U/ml RNasin at 40 °C and then washed four times in wash buffer, PBS, 0.1% (vol/vol) Tween 20, and 4 U/ml RNasin (Promega, Madison, WI, USA). Samples were washed and endogenous biotin was blocked using Avidin/Biotin blocking kit (Vector laboratories, Burlingame, CA, USA) according to the manufacture instructions. Slides were incubated with a biotin labeled DNA probe at a concentration of 10 nM at 37 °C in PBS, 1× SSC, 0.1% Tween 20, 100 μg/ml salmon sperm DNA, 100 μg/ml yeast RNA. After washing, samples were incubated with 1:100 diluted streptavidin-HRP (Thermo Fisher Scientific, Waltham, MA, USA) in PBS, 60 min at room temperature followed by washing. Samples were labeled using ImmPACT-DAB substrate, counter stained using hematoxylin QS and imbedded in VectaMount AQ mounting medium (Vector laboratories, Burlingame, CA, USA) according to the manufacture instructions.

### Data presentation

Figures arranged in Adobe Illustrator 2020.

## Competing interests

The Rockefeller University has filed a provisional patent application in connection with this work on which D.F.R. and M.C.N. are inventors (US patent 63/021,387). R.E.S. is on the scientific advisory board of Miromatrix Inc and is a consultant and speaker for Alnylam Inc. S.M. has served as a consultant for Takeda Pharmaceuticals, Morphic and Glaxo Smith Kline. Z.Z. received seed instruments and sponsored travel from ET Healthcare.

## Data availability statement

Data are provided in SI Tables 1-8. The raw sequencing data and computer scripts associated with Figure 2 have been deposited at Github (https://github.com/stratust/igpipeline). This study also uses data from “A Public Database of Memory and Naive B-Cell Receptor Sequences” (https://doi.org/10.5061/dryad.35ks2), PDB (6VYB and 6NB6) and from “High frequency of shared clonotypes in human B cell receptor repertoires” (https://doi.org/10.1038/s41586-019-0934-8).

## Code availability statement

Computer code to process the antibody sequences is available at GitHub (https://github.com/stratust/igpipeline).

## Acknowledgements

We thank all study participants who devoted time to our research; Drs. Barry Coller and Sarah Schlesinger, the Rockefeller University Hospital Clinical Research Support Office and nursing staff. Mayu Okawa Frank and Robert B. Darnell for SARS-CoV-2 saliva PCR testing. Charles M. Rice and all members of the M.C.N. laboratory for helpful discussions and Maša Jankovic for laboratory support. This work was supported by NIH grant P01-AI138398-S1 (M.C.N. and P.J.B.) and 2U19AI111825 (M.C.N.).; the Caltech Merkin Institute for Translational Research and P50 AI150464-13 (P.J.B.), George Mason University Fast Grant (D.F.R. and P.J.B.) and the European ATAC consortium EC 101003650 (D.F.R.); R37-AI64003 to P.D.B.; R01AI78788 to T.H.; NCI R01CA234614, NIAID 2R01AI107301, NIDDK R01DK121072 and 1RO3DK117252 to R.E.S., NIH NIDDK R01 DK123749 01S1 to S.M. We thank Dr. Jost Vielmetter and the Protein Expression Center in the Beckman Institute at Caltech for expression assistance. C.O.B. is supported by the HHMI Hanna Gray and Burroughs Wellcome PDEP fellowships. R.E.S. is a Irma Hirschl Trust Research Award Scholar. M.T. is supported by the Digestive Disease Research Foundation (DDRF). C.G. was supported by the Robert S. Wennett Post-Doctoral Fellowship, in part by the National Center for Advancing Translational Sciences (National Institutes of Health Clinical and Translational Science Award program, grant UL1 TR001866), and by the Shapiro-Silverberg Fund for the Advancement of Translational Research. P.D.B. and M.C.N. are Howard Hughes Medical Institute Investigators.

## Author Contributions

C.G., P.D.B., P.J.B., T.H., S.M. and M.C.N. conceived, designed and analyzed the experiments. D.F.R., M.Caskey. and C.G. designed clinical protocols. Z.W., J.C.C.L., F.M., S.F., M.T., A.C., M.J., D.S.B., F.S., Y.W., T.H., P.M., G.B., C.V., C.O.B., K.G., D.J., J.Y., G.M.D., Y.B., R.E.S. and Z.Z. carried out experiments. A.G. and M.Cipolla produced antibodies. A.H., D.S.B., M.Turroja, K.G.M., M.Tankelevich, C.G. and M.Caskey recruited participants and executed clinical protocols. I.S., R.P, J.D. and C.U.O. processed clinical samples. T.Y.O. and V.R. performed bioinformatic analysis. C.G, P.D.B., P.J.B., T.H., S.M. and M.C.N. wrote the manuscript with input from all co-authors.

## Extended Data Legends

**Extended Data Fig. 1:**
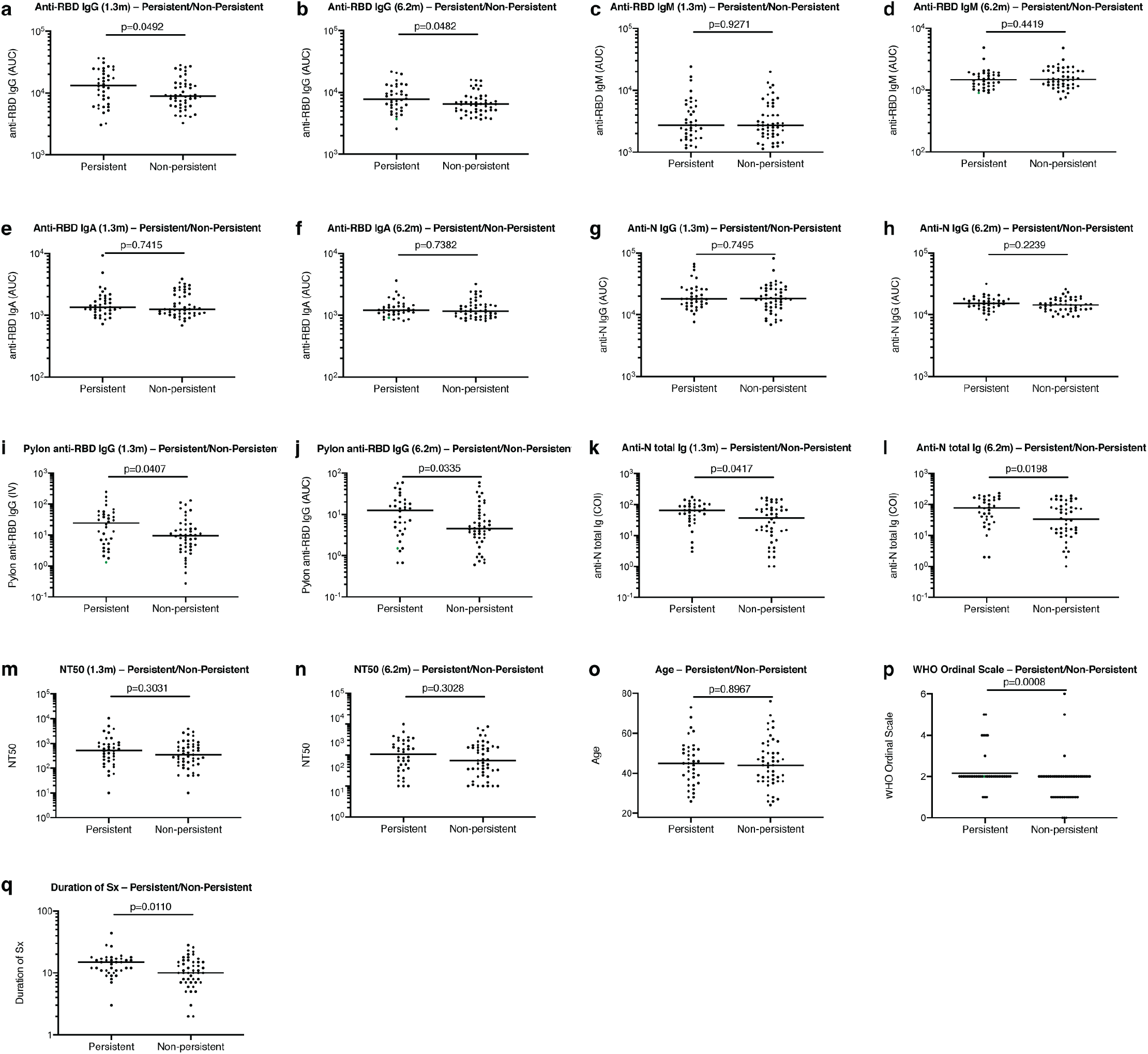
Clinical correlates of plasma antibody titres. **a**, Normalized AUC anti-RBD IgG titres at 1.3 months for participants with (n=38) or without (n=49) persistent post-acute symptoms. **b**, Normalized AUC anti-RBD IgG titres at 6.2 months for participants with (n=38) or without (n=49) persistent post-acute symptoms. **c**, Normalized AUC anti-RBD IgM titres at 1.3 months for participants with (n=38) or without (n=49) persistent post-acute symptoms. **d**, Normalized AUC anti-RBD IgM titres at 6.2 months for participants with (n=38) or without (n=49) persistent post-acute symptoms. **e**, Normalized AUC anti-RBD IgA titres at 1.3 months for participants with (n=38) or without (n=49) persistent post-acute symptoms. **f**, Normalized AUC anti-RBD IgA titres at 6.2 months for participants with (n=38) or without (n=49) persistent post-acute symptoms. **g**, Normalized AUC anti-N IgG titres at 1.3 months for participants with (n=38) or without (n=49) persistent post-acute symptoms. **h**, Normalized AUC anti-N IgG titres at 6.2 months for participants with (n=38) or without (n=49) persistent post-acute symptoms. **i**, IV values of anti-RBD IgG titres at 1.3 months for participants with (n=38) or without (n=49) persistent post-acute symptoms. **j**, IV values of anti-RBD IgG titres at 6.2 months for participants with (n=38) or without (n=49) persistent post-acute symptoms. **k**, COI values of anti-N total Ig titres at 1.3 months for participants with (n=38) or without (n=49) persistent post-acute symptoms. **l**, COI values of anti-N total Ig titres at 6.2 months for participants with (n=38) or without (n=49) persistent post-acute symptoms. **m**, NT50 values at 1.3 months for participants with (n=38) or without (n=49) persistent post-acute symptoms. **n**, NT50 values at 6.2 months for participants with (n=38) or without (n=49) persistent post-acute symptoms. **o**, Age in years for participants with (n=38) or without (n=49) persistent post-acute symptoms. **p**, Severity of acute infection as assessed by the WHO Ordinal Clinical Progression/Improvement Scale for participants with (n=38) or without (n=49) persistent post-acute symptoms. **q**, Duration of Symptoms during acute infection for participants with (n=38) or without (n=49) persistent post-acute symptoms. Horizontal bars indicate median values. Statistical significance was determined using two-tailed Mann–Whitney U-tests.

**Extended Data Fig. 2:**
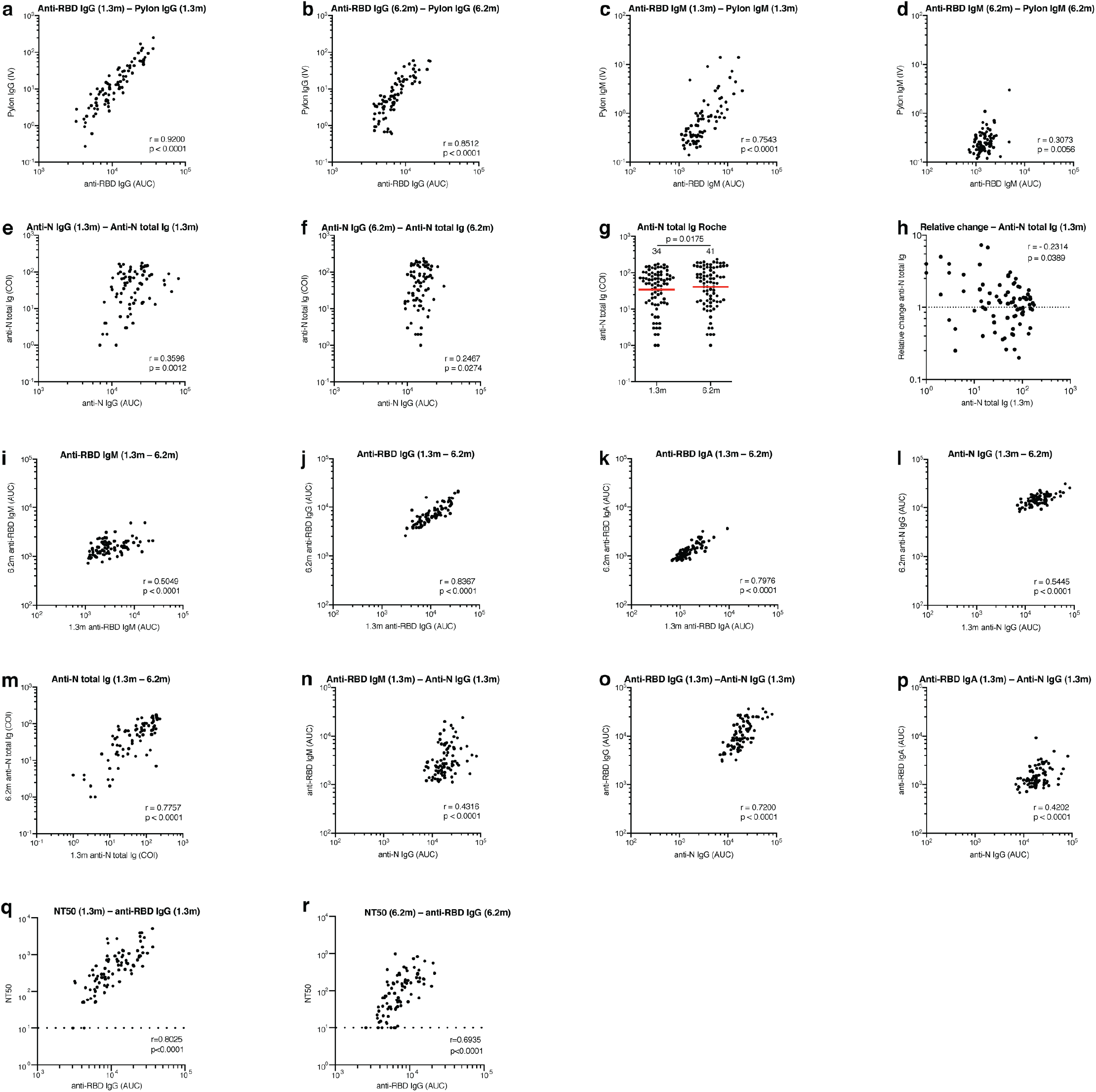
Correlations of plasma antibody measurements. **a,** Normalized AUC for IgG anti-RBD plotted against Pylon IgG anti-RBD index values at 1.3 months. **b,** Normalized AUC for IgG anti-RBD plotted against Pylon IgG anti-RBD index values at 6.2 months. **c,** Normalized AUC for IgM anti-RBD plotted against Pylon IgM anti-RBD index values at 1.3 months. **d,** Normalized AUC for IgM anti-RBD plotted against Pylon IgM anti-RBD index values at 6.2 months. **e**, Normalized AUC for IgG anti-N plotted against Roche COI values for anti-N total Ig titres at 1.3 months. **f,** Normalized AUC for IgG anti-N plotted against Roche COI values for anti-N total Ig titres at 6.2 months. **g**, Anti-N total Ig Cut-off Index (COI) values for 80 individuals at the initial 1.3 and 6.2 month follow-up visit. **h**, Relative change in anti-N total Ig levels between 1.3 and 6.2 months plotted against the anti-N total Ig levels at 1.3 months. **i,** Normalized AUC for IgM anti-RBD at 6.2 months plotted against IgM anti-RBD at 1.3 months. **j**, Normalized AUC for IgG anti-RBD at 6.2 months plotted against IgG anti-RBD at 1.3 months. **k**, Normalized AUC for IgA anti-RBD at 6.2 months plotted against IgA anti-RBD at 1.3 months. **l**, Normalized AUC for IgG anti-N at 6.2 months plotted against IgG anti-N at 1.3 months. **m**, COI values for anti-N total Ig titres at 6.2 months plotted against anti-N total Ig titres at 1.3 months. **n**, Anti-RBD IgM titres at 1.3 months plotted against anti-N IgG titres at 1.3 months. **o,** Anti-RBD IgG titres at 1.3 months plotted against anti-N IgG titres at 1.3 months. **p,** Anti-RBD IgA titres at 1.3 months plotted against anti-N IgG titres at 1.3 months. **q**, NT50 values at 1.3 months plotted against anti-RBD IgG titres at 1.3 months. **r**, NT50 values at 6.2 months plotted against anti-RBD IgG titres at 6.2 months. The *r* and *p* values were determined by two-tailed Spearman’s correlations.

**Extended Data Fig. 3:**
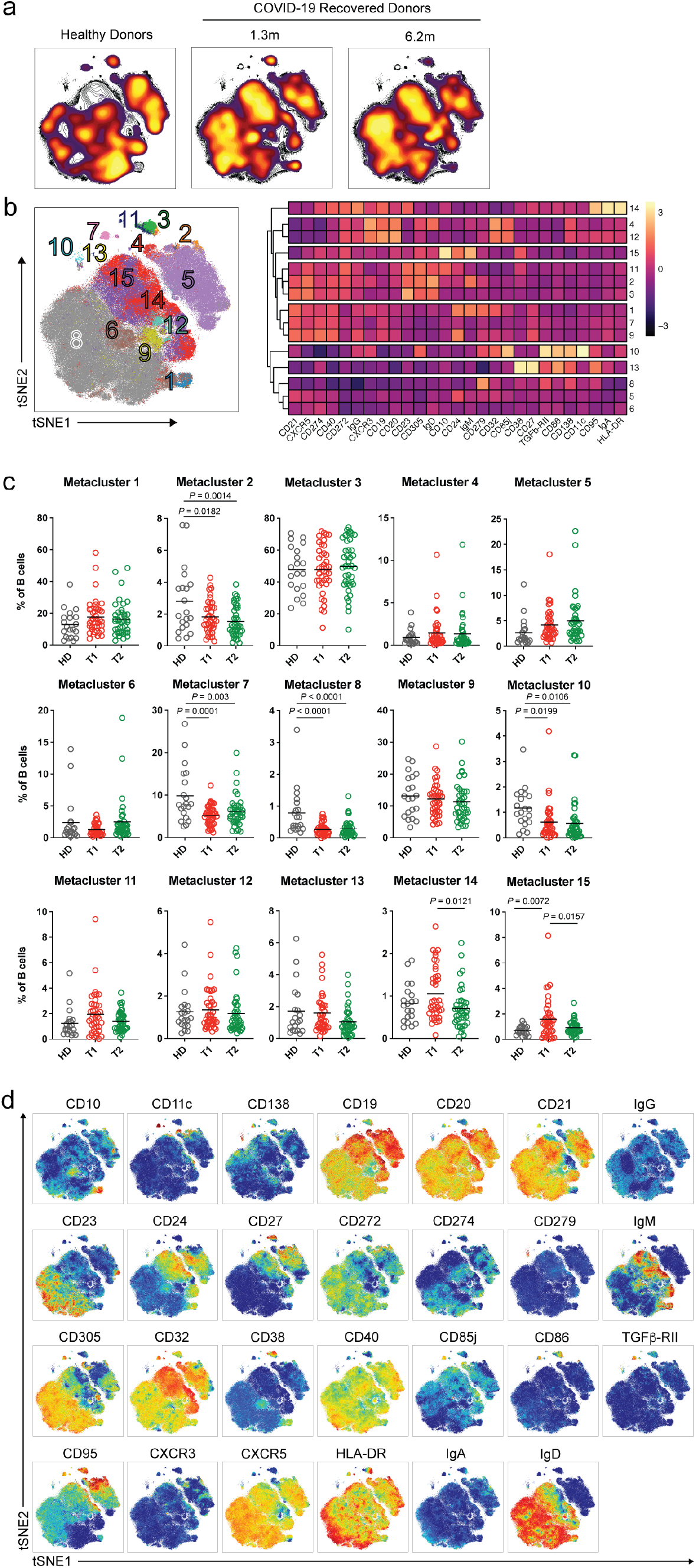
Persistent longitudinal changes in the phenotypic landscape of B cells in individuals recovered from COVID-19. **a**, Global viSNE projection of pooled B cells for all participants pooled shown in background contour plots, with overlaid projections of concatenated controls, convalescent participants 1.3 and 6.1 months respectively. **b**, viSNE projection of pooled B cells for all participants of B cell clusters identified by FlowSOM clustering. Column-scaled z-scores of median fluorescence intensity (MFI) as indicated by cluster and marker. **c**, Frequency of B cells from each group in FlowSOM clusters indicated. Each circle represents an individual control individual (n=20) (gray), convalescent participant at 1.3 months post-infection (n=41) (red), or convalescent participant at 6.2 months post-infection (n=41) (green). Horizontal bars indicate mean values. Significance determined by two-tailed paired t-test for comparisons between time points within individuals and two-tailed unpaired t-test for comparison between controls and convalescent individuals. **d**, Individual viSNE projections of indicated protein expression.

**Extended Data Fig. 4:**
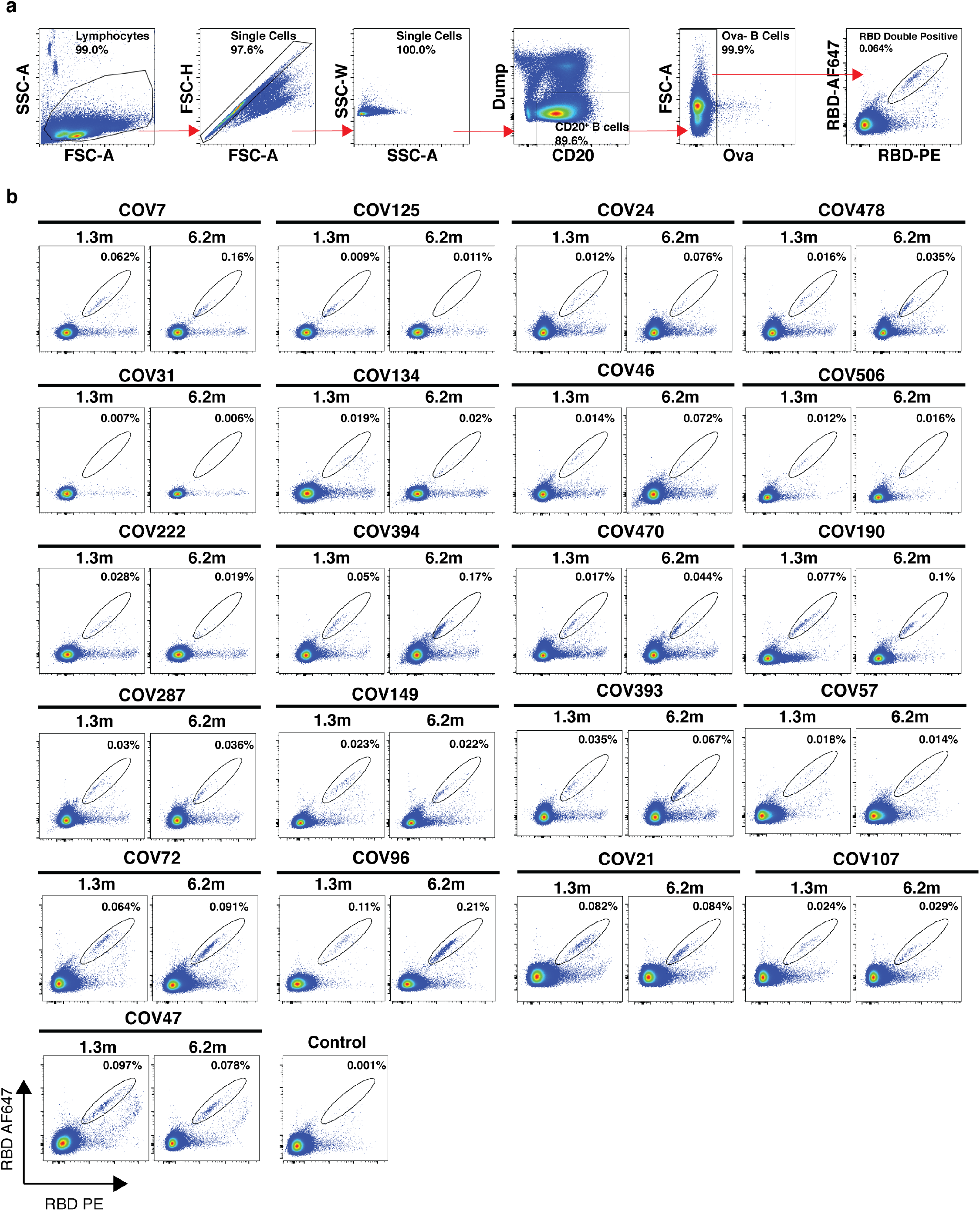
Flow cytometry. **a,** Gating strategy used for cell sorting. Gating was on singlets that were CD20+ and CD3−CD8−CD16−Ova−. Sorted cells were RBD-PE+ and RBD-AF647+. **b,** Flow cytometry showing the percentage of RBD-double positive memory B cells from month 1.3 or month 6 post-infection in 21 randomly selected participants.

**Extended Data Fig. 5:**
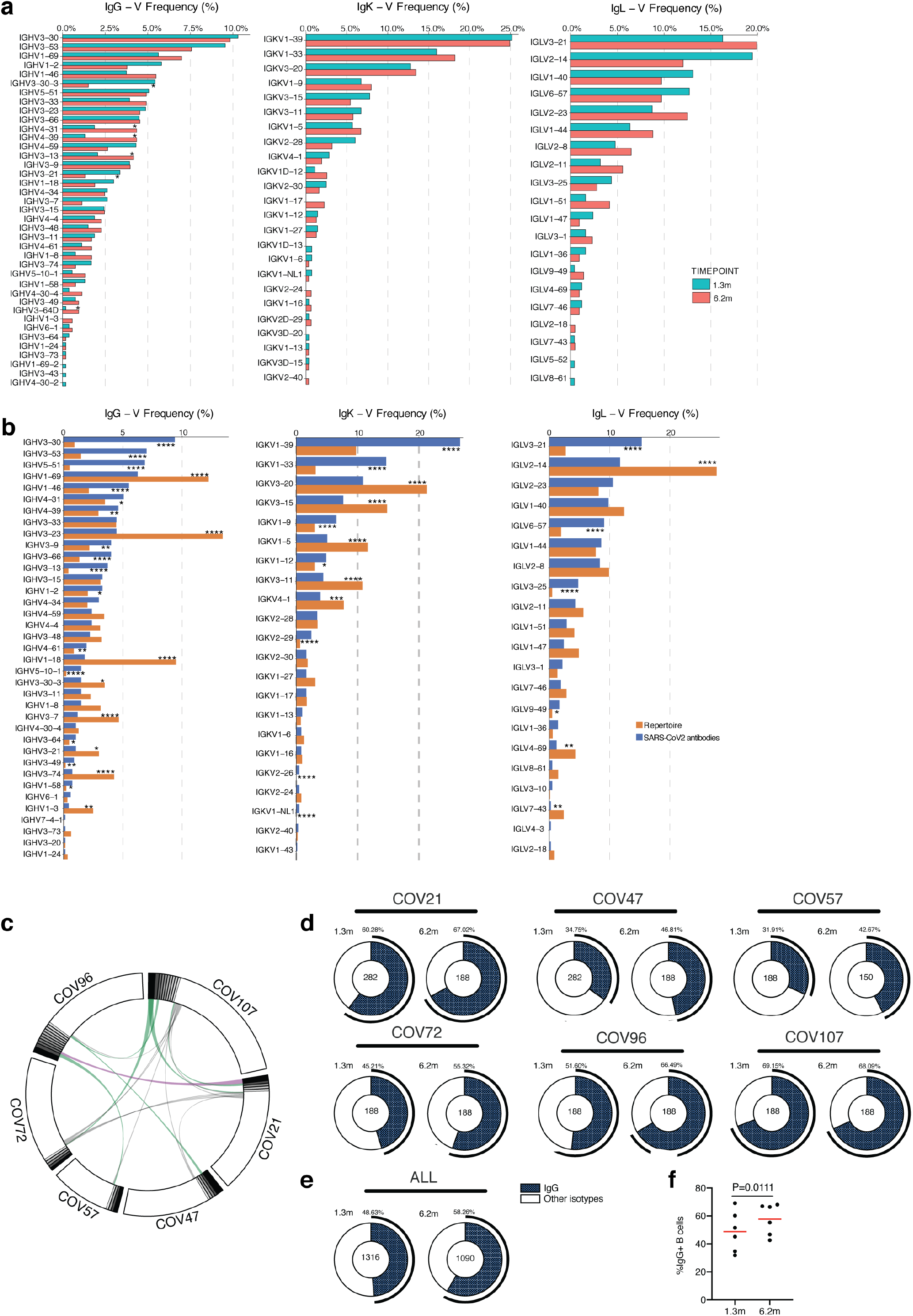
Frequency distributions of human V genes. **a,** Two-sided binomial tests were used to compare the frequency distributions of human V genes of anti-SARS-CoV-2 antibodies from donors at 1.3 months to 6.2 months ^1^ (* p < 0.05, ** p < 0.01 *** p < 0.001, **** = p < 0.0001). **b,** Two-sided binomial tests were used to compare the frequency distributions of human V genes of anti-SARS-CoV-2 antibodies from this study to sequence from *C. Soto et al*. ^54^ (* p < 0.05, ** p < 0.01, *** p < 0.001, **** = p < 0.0001). **c**, Sequences from all six individuals with clonal relationships depicted as circos plots as in Figure 2d. Interconnecting lines indicate the relationship between antibodies that share V and J gene segment sequences at both IGH and IGL. Purple, green and grey lines connect related clones, clones and singles, and singles to each other, respectively. **d,** For each participant, the number of IgG heavy chain sequences (black) analyzed at month 1.3 (left panel) or month 6.2 post-infection (right panel). The number in the inner circle indicates the number of cells that was sorted for each individual denoted above the circle. **e,** The same as d but showing combined data for all six participants. **f,** Comparison of the percentage of IgG positive B cells from all six individuals at month 1.3 or month 6.2 post-infection. The horizontal bars indicate the mean. Statistical significance was determined using two-tailed t-test.

**Extended Data Fig. 6:**
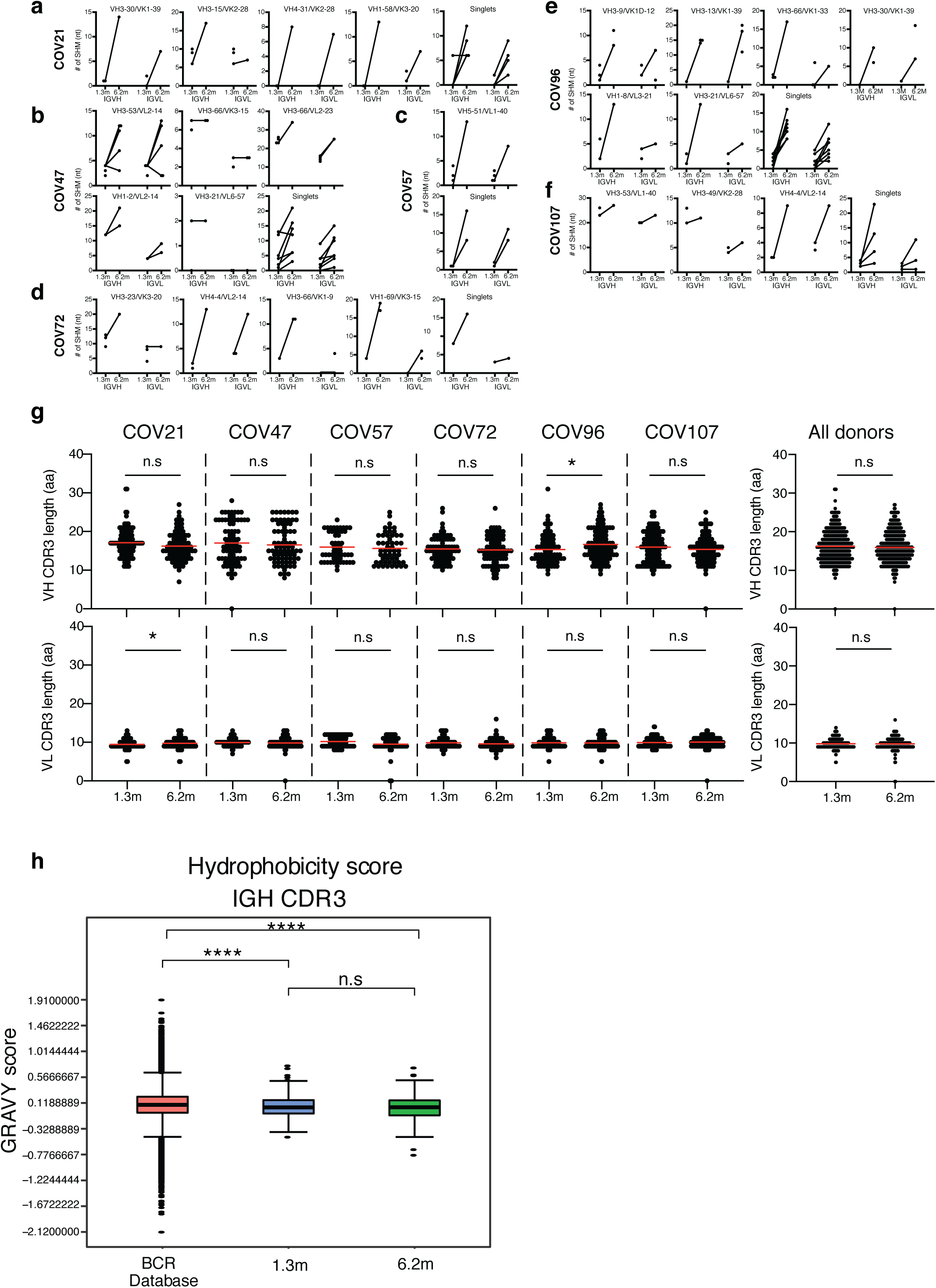
Analysis of antibody somatic hypermutation of persisting clones, CDR3 length and hydrophobicity. **a-f**, Number of somatic nucleotide mutations in both the IGVH and IGVL of persisting clones found at month 1.3 and month 6.2 time points in six participants (a) COV21, (b) COV47, (c) COV57, (d) COV72, (e) COV96, and (f) COV107. The VH and VL gene usage of each clonal expansion is indicated above the graphs, or are indicated as “Singlets” if the persisting sequence was isolated only once at both time points. Connecting line indicates the SHM of the clonal pairs that were expressed as a recombinant mAbs. **g,** For each individual, the number of the amino acid length of the CDR3s at the IGVH and IGVL is shown. The horizontal bars indicate the mean. The number of antibody sequences (IGVH and IGVL) evaluated for each participant are n = 90 (COV21), n = 78 (COV47), n = 53 (COV57), n = 87 (COV72), n = 104 (COV96), n = 120 (COV107). Right panel show all antibodies combined (n = 532 for both IGVH and IGVL). Statistical significance was determined using two-tailed Mann–Whitney U-tests and horizontal bars indicate median values (n.s.=non-significant, * p < 0.05, ** p < 0.01, *** p < 0.001, **** = p < 0.0001). **h,** Distribution of the hydrophobicity GRAVY scores at the IGH CDR3 in 532 antibody sequences from this study compared to a public database (see Methods for statistical analysis). The box limits are at the lower and upper quartiles, the center line indicates the median, the whiskers are 1.5× interquartile range and the dots represent outliers. Statistical significance was determined using two-tailed Wilcoxon matched-pairs signed rank test (n.s.=non-significant, * p < 0.05, ** p < 0.01, *** p < 0.001, **** = p < 0.0001).

**Extended Data Fig. 7:**
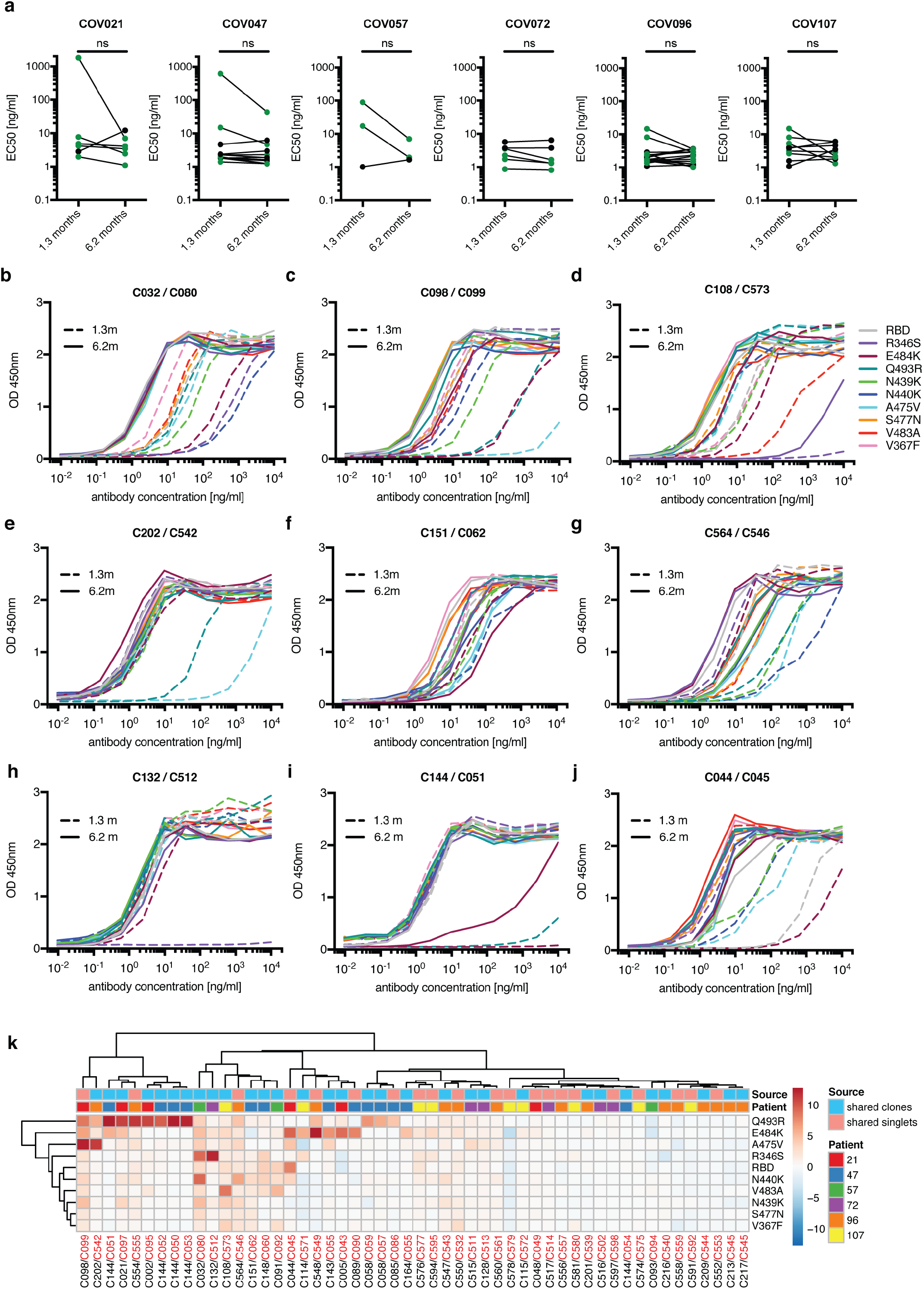
ELISA of wt/mutant RBD for mAbs. **a,** EC50 values for binding to wild type RBD of shared singlets and shared clones of mAbs obtained at the initial 1.3 and 6.2 months follow-up visit, divided by participant (n = 6 (COV21), n = 13 (COV47), n = 3 (COV57), n = 6 (COV72), n = 15 (COV96), n = 9 (COV107). Lines connect shared singlets/clones. mAbs with improved EC50 at 6.2 months follow-up visit are highlighted in green, remaining mAbs are shown in black. Statistical significance was determined using two-tailed Wilcoxon matched-pairs signed rank test. **b-j,** Graphs show ELISA binding curves for different antibodies obtained at 1.3 months (dashed lines) and their clonal relatives obtained after 6.2 months (solid lines) binding to wild type, R346S, E484K, Q493R, N439K, N440K, A475V, S477N, V483A and V367F RBDs (colors as indicated). Antibody IDs of pairs as indicated on top of panels (1.3m / 6.2m). **k,** Heat map shows log2 relative fold change in EC_50_ against the indicated RBD mutants for 52 antibody clonal pairs obtained at 1.3 (black) and 6.2 months (red). The clonal and participant origin for each antibody pair is indicated above.

**Extended Data Fig. 8:**
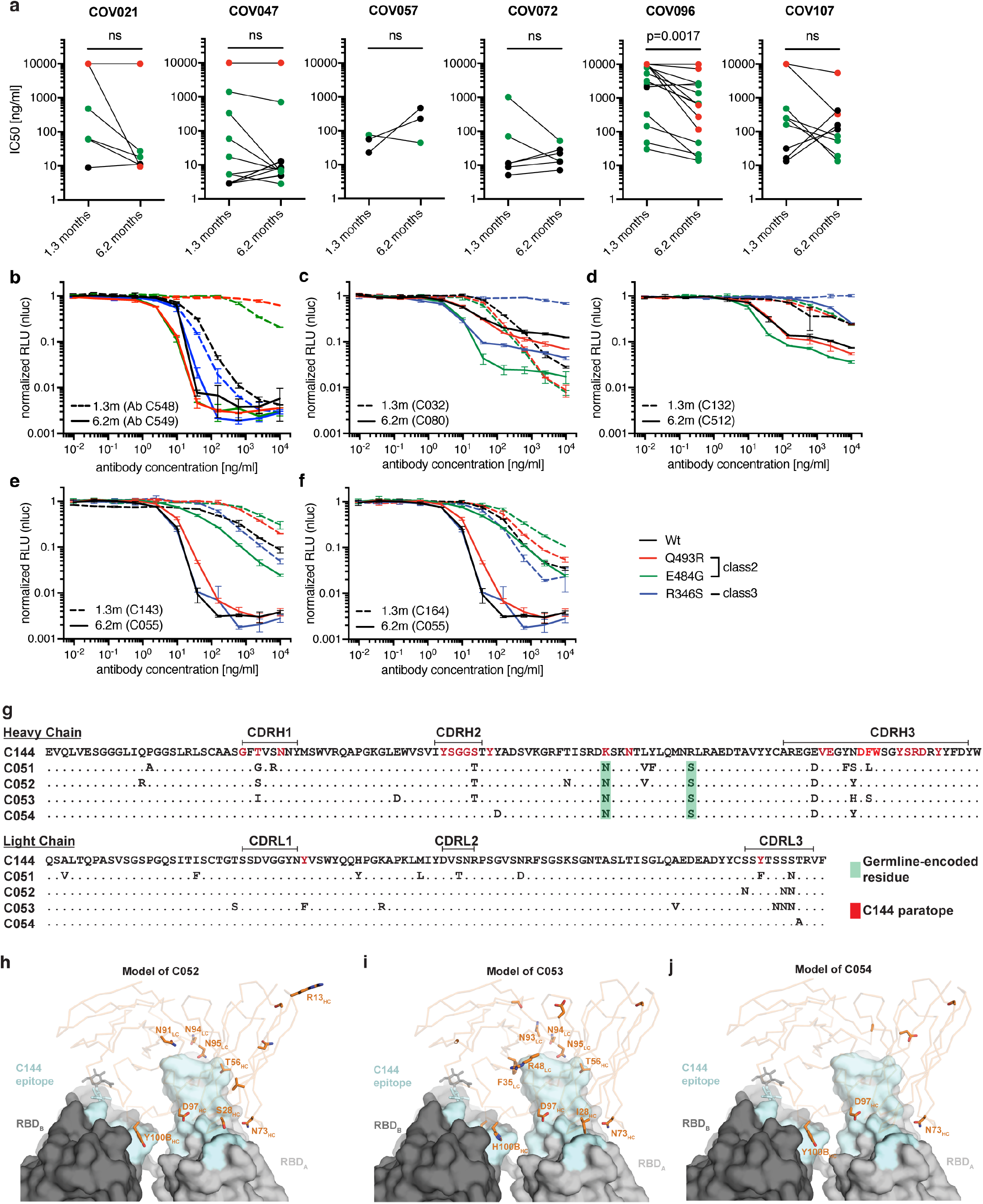
Neutralization of wt/mutant RBDs, C51 alignment and binding projection. **a,** IC50 values of shared singlets and shared clones of mAbs obtained at the initial 1.3-and 6.2-months follow-up visit, divided by participant (n = 6 (COV21), n = 13 (COV47), n = 3 (COV57), n = 6 (COV72), n = 15 (COV96), n = 9 (COV107). Lines connect shared singlets/clones. mAbs with undetectable IC50 at 1.3 months are plotted at 10 µg/ml and are highlighted in red, mAbs with improved IC50 at 6.2 months follow-up visit are highlighted in green, remaining mAbs are shown in black. Statistical significance was determined using two-tailed Wilcoxon matched-pairs signed rank test. **b-f,** The normalized relative luminescence values for cell lysates of 293TAce2 cells 48 hpi with SARS-CoV-2 pseudovirus harboring wt RBD or RBD-mutants (wt, Q493R, E484G and R346S RBD shown in black, red, green and blue, respectively) in the presence of increasing concentrations of mAbs obtained at the 1.3 months initial visit (1.3m, dashed lines) and their shared clones/singlets at the 6.2 follow-up visit (6.2m, continuous lines). Antibody IDs as indicated. **g,** VH and VL amino acid sequence alignment of C144 and derivative antibodies C051, C052, C053 and C054. Germline-encoded residues are highlighted in green. Residues in the proximity of RBD-binding C144 paratope are highlighted in red. **h-j**, Surface representation of two adjacent “down” RBDs (RBD_A_ and RBD_B_) on a spike trimer with the C144 epitope on the RBDs highlighted in cyan and positions of amino acid mutations that accumulated in **h,** C052. **i**, C053 and **j**, C054 compared to the parent antibody C144 highlighted as stick side chains on a Cα atom representation. The C052, C053 and C054 interactions with two RBDs was modeled based on a cryo-EM structure of C144 Fab bound to spike trimer ^17^.

**Extended Data Fig. 9:**
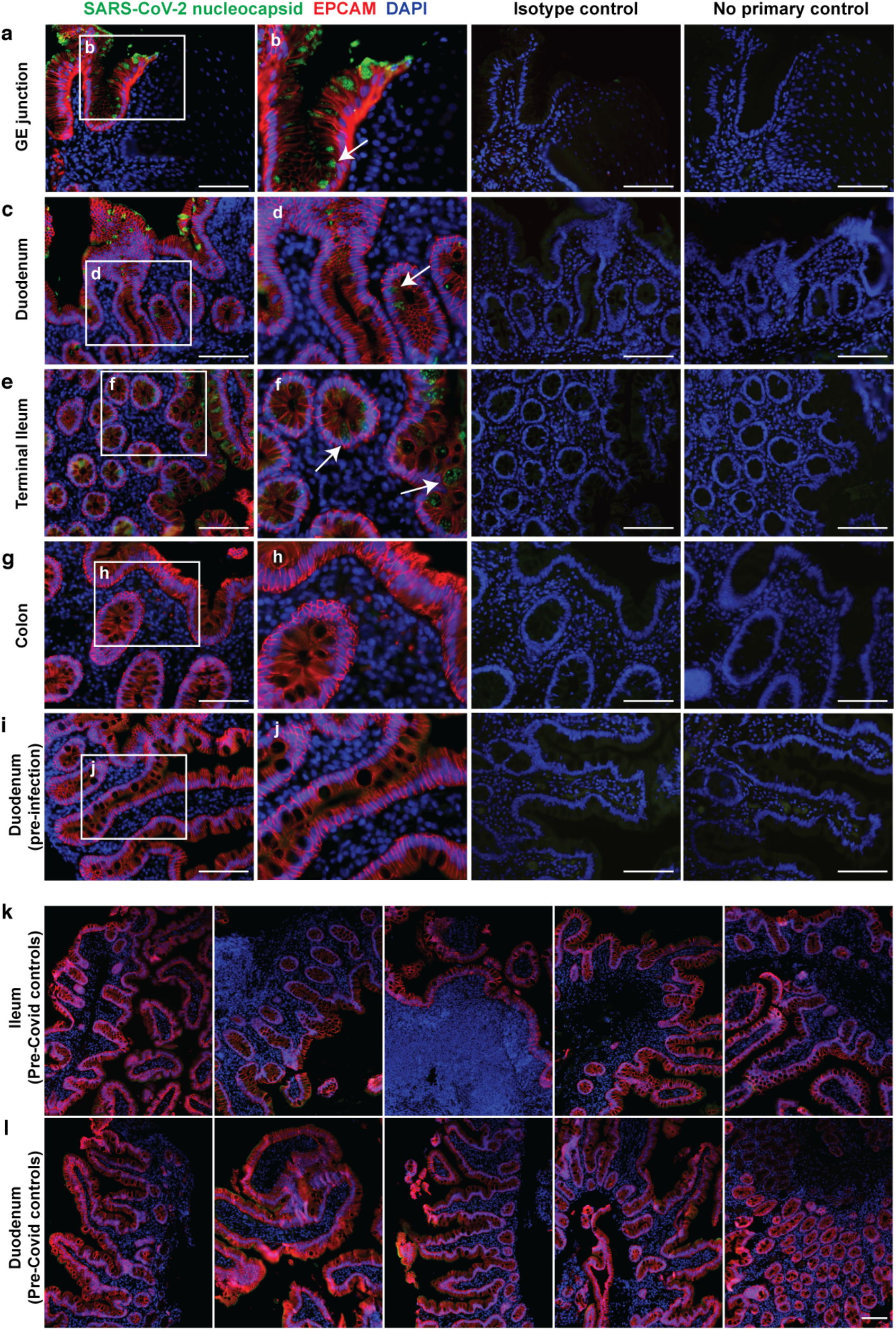
SARS-CoV-2 antigen in human enterocytes in the gastrointestinal tract 3 months post COVID-19 and Pre-COVID-19 historic control individuals without detectable SARS-CoV-2 antigen. **a-j**, Immunofluorescence images of human gastrointestinal tissue are shown. Staining is for EPCAM (red), DAPI (blue) and SARS-CoV-2 nucleocapsid (green) Samples are derived from intestinal biopsies in the gastrointestinal tract as indicated (a-j). (a-h) are biopsies from participant CGI-088 (Supplementary Table 7) taken 92 days post COVID-19 symptom onset. (i-j) is a biopsy 27 months prior to COVID-19 symptom onset from the same participant CGI-088. Arrows indicate enterocytes with detectable SARS-CoV-2 antigen. Isotype and no primary controls for each tissue are shown in the right two columns. White scale bar corresponds to 100 μm. **k-l**, Immunofluorescence images of biopsy samples in the gastrointestinal tract (k-ileal, l-duodenal) obtained from 10 different Pre-COVID-19 historic control individuals between January 2018 and October 2019 are shown. Staining is for EPCAM (red), DAPI (blue) and SARS-CoV-2 nucleocapsid (green). White scale bar corresponds to 100 μm. All experiments were repeated independently at least twice with similar results.

**Extended Data Fig. 10:**
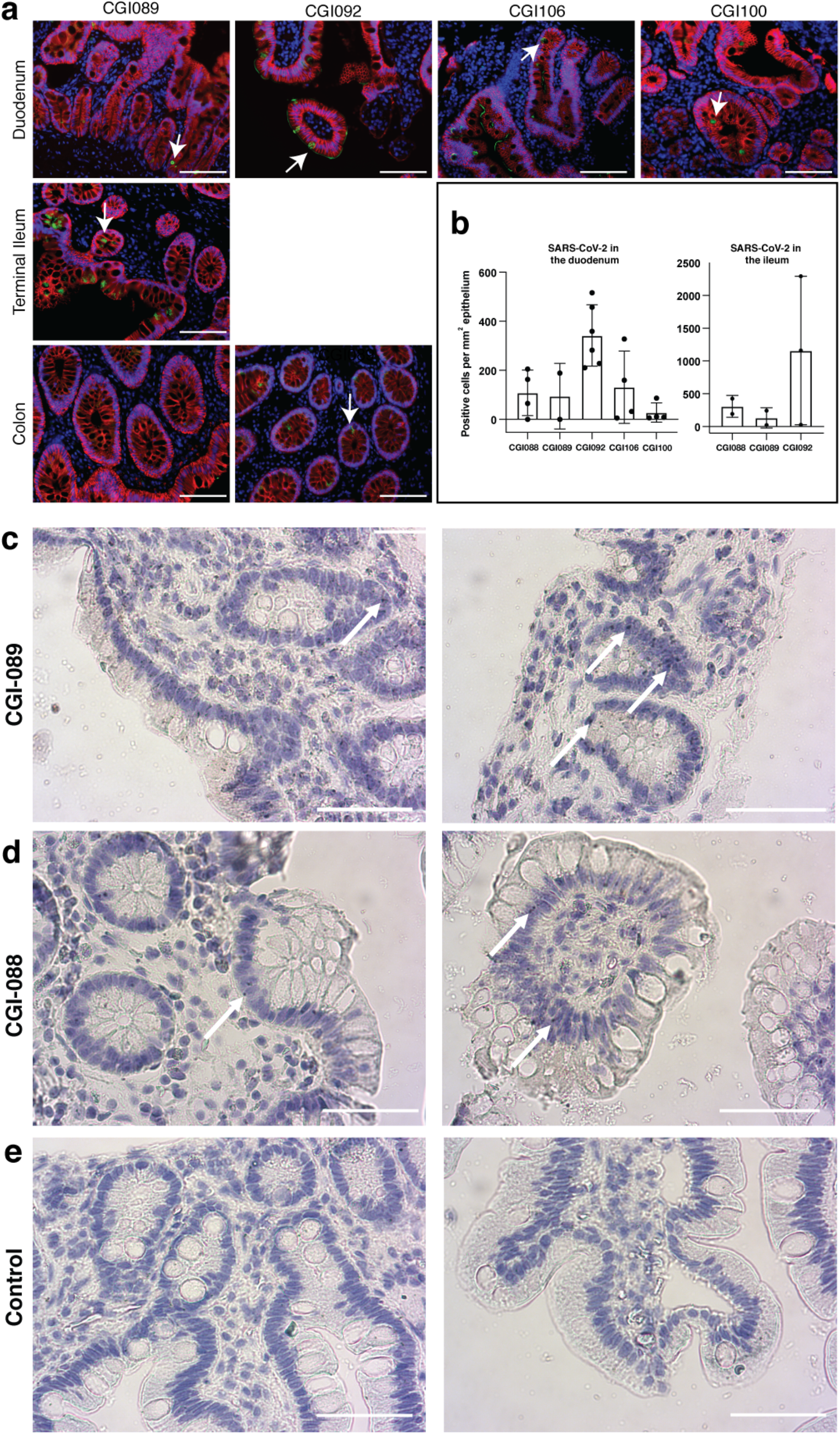
SARS-CoV-2 antigen and RNA is detectable in different intestinal segments in multiple COVID-19 convalescent individuals. **a**, Immunofluorescence (IF) images of biopsy samples in the gastrointestinal tract in different individuals are shown. Staining is for EPCAM (red), DAPI (blue) and SARS-CoV-2 nucleocapsid (green). Samples are derived from intestinal biopsies from 4 participants (CGI089, CGI092, CGI100 and CGI106) taken at least 3 months after COVID-19 infection. Arrows indicate enterocytes with detectable SARS-CoV-2 antigen. White scale bar corresponds to 100 μm. The experiments were repeated independently at least twice with similar results. **b**, Quantification of SARS-CoV-2 positive cells by immunofluorescence. The number of cells staining positive for the nucleocapsid protein (N) of SARS-CoV-2 per mm^2^ of intestinal epithelium is shown. The graphs show biopsy samples from the indicated individuals of the duodenum (left) and terminal ileum (right), respectively. Black dots represent the number of available biopsy specimen for each individual from the respective intestinal segment (CGI-088 n=4 duodenal/n=2 ileal, CGI-089 n=2 duodenal/n=2 ileal, CGI-092 n=6 duodenal/n=3 ileal, CGI-106 n=4, CGI-100 n=4). Boxes represent median values and whiskers the 95 % CI. **c-d,** SARS-CoV-2 viral RNA was visualized in intestinal biopsies of participant CGI-089 (c) and CGI-088 (d) using in situ hybridization. SARS-CoV-2 genomic RNA (black) and Hematoxylin and Eosin staining by smFISH/IHC technique in the duodenum (left) or terminal ileum (right). Arrows indicate enterocytes with detectable SARS-CoV-2 RNA. **e**, Pre-COVID-19 historic control individuals show no detectable SARS-CoV-2 viral RNA in duodenal (left) or ileal (right) biopsies. White scale bar corresponds to 100 μm. The experiments were repeated independently three times with similar results.

## Supplementary Tables

**Supplementary Table 1: Cohort summary is provided as a separate Excel file.**

**Supplementary Table 2: Individual participant characteristics is provided as a separate Excel file.**

**Supplementary Table 3: Antibody sequences from 1.3 and 6.2 month time point is provided as a separate Excel file.**

**Supplementary Table 4: Sequences of the cloned monoclonal antibodies is provided as a separate Excel file.**

**Supplementary Table 5: Half maximal effective concentrations (EC50s) of the monoclonal antibodies is provided as a separate Excel file.**

**Supplementary Table 6: Inhibitory concentrations of the monoclonal antibodies is provided as a separate Excel file.**

**Supplementary Table 7: Gastrointestinal biopsies cohort characteristics is provided as a separate Excel file.**

**Supplementary Table 8: Proximity ligation and smFISH probe sequences is provided as a separate Excel file.**

